# The role of mitotype variation and positive epistasis in trait differences between *Saccharomyces* species

**DOI:** 10.1101/2025.05.07.652752

**Authors:** Jun-Ting Johnson Wang, Ping Ling Priscilla Ng, Maceo E. Powers, Catherine H. Rha, Rachel B. Brem

## Abstract

Many traits of interest in biology evolved long ago and are fixed in a particular species, distinguishing it from other sister taxa. Elucidating the mechanisms underlying such divergences across reproductive barriers has been a key challenge for evolutionary biologists. The yeast *Saccharomyces cerevisiae* is unique among its relatives for its ability to thrive at high temperature. The genetic determinants of the trait remain incompletely understood, and we sought to understand the role in its architecture of species variation in mitochondrial DNA. We used mitochondrial transgenesis to show that *S. cerevisiae* mitotypes were sufficient for a partial boost to thermotolerance and respiration in the *S. paradoxus* background. These mitochondrial alleles worked best when the background also harbored a pro-thermotolerance nuclear genotype, attesting to positive epistasis between the two genomes. The benefits of *S. cerevisiae* alleles in terms of respiration and growth at high temperature came at the cost of worse performance in cooler conditions. Together, our results establish this system as a case in which mitoalleles drive fitness benefits in a manner compatible with, and fostered by, the nuclear genome.

## Introduction

A central aim in evolutionary genetics is to figure out how organisms acquire new traits. In the study of within-species variation, a rich literature has provided insights into the mechanisms of local adaptation in the wild (Elena 2017; Kraemer and Boynton 2017; Sork 2017; Rees et al. 2020; Bazzicalupo 2022). Yet many characters of basic and applied interest evolved long ago and are fixed in a particular species, distinguishing it from unaffected relatives. Against a backdrop of years of genomic and experimental pursuit (Allen Orr 2001; Nikolov and Tsiantis 2015; Weiss and Brem 2019), the genetic principles governing trait divergence over long timescales remain poorly understood, with the widest knowledge gap in the case of polygenic architectures. Indeed, an adaptive trait that arose long ago, and ultimately fixed in a given species, could have undergone millions of years of refinement, and could have a genetic architecture governed by principles quite different from recently-arisen intra-species trait polymorphism that is the focus of much of the field. Shedding light on these mechanisms, by dissecting the genetics of adaptations between isolated taxa, remains a key challenge.

Sequence divergence in mitotype, the DNA of the mitochondrial organelle, have emerged as an important driver of naturally varying traits. Mitotype variants have been directly implicated in trait diversity, both independently and in concert with nuclear genomes, in a rich literature focused on polymorphism within species (Melvin and Ballard 2006; Tranah 2011; Houtkooper et al. 2013; Sullivan and Chandel 2014; Mossman et al. 2016 Nov 14; Mossman et al. 2016; Patel et al. 2016; Camus et al. 2017; Camus and Dowling 2018; Mossman et al. 2019; Sun et al. 2019; Camus et al. 2020; Salminen and Vale 2020; Anderson et al. 2022; Cao et al. 2022; Meng et al. 2022; Oppong et al. 2022; Quéméneur et al. 2022; Parra et al. 2023; Moran et al. 2024). Though small in length, the mitochondrial genome has the potential for an outsize influence on trait variation, given its increased mutation rate and large mutational target of the multi-copy mitochondrial genome relative to nuclear chromosomes (Pakendorf and Stoneking 2005; Chou and Leu 2015; Ferreira and Rodriguez 2024).

*Saccharomyces* budding yeasts diverged from a common ancestor ~20 million years ago (Kellis et al. 2003) and have served for decades as a model for the study of interspecific trait variation, including in metabolism, genome content, cell cycle, and reproductive isolation (Hou et al. 2014; Roop et al. 2016; Bozdag et al. 2021; Fredericks et al. 2021; Lupo et al. 2021; Marsit et al. 2021; Smukowski Heil et al. 2021; Swamy et al. 2022; Crandall et al. 2023; Forejt and Jansa 2023; Peris et al. 2023). A long-standing subset of this field has focused on species-unique traits driven by mitochondrial genomes (Sulo et al. 2003; Wolff et al. 2014; De Chiara et al. 2020; Hewitt et al. 2020; Hénault et al. 2022), with particularly detailed mechanistic validation in studies of speciation (Lee et al. 2008; Chou et al. 2010; Mário et al. 2015; Jhuang et al. 2017) (complementing studies of mitotype polymorphism within yeast species (Wolters et al. 2015; Leducq et al. 2017; Wolters et al. 2018; Vijayraghavan et al. 2019; Vijayraghavan et al. 2019; Nguyen et al. 2020)). A key model trait in the *Saccharomyces* system is temperature preference, as the best-studied yeast species, *S. cerevisiae*, is unique within the *Saccharomyces* clade for its ability to thrive at high temperature (Sweeney et al. 2004; Gonçalves et al. 2011; Salvadó et al. 2011), whereas others in the clade exhibit cryotolerance (Paget et al. 2014; Pinto et al. 2025). Species-level variation in these traits has been inaccessible to analysis by classic methods like GWAS or linkage mapping, which rely on recombinants in fertile crosses. For this reason, mechanisms by which nature built thermotolerance in *S. cerevisiae* from a thermosensitive ancestor were difficult to access for the classic literature. But the system represents a testbed for alternative approaches for genetic dissection across species boundaries, which have fostered insight into the architectures of species differences in thermotolerance and their adaptive history (Weiss et al. 2018; AlZaben et al. 2021; Walunjkar et al. 2025) and cold tolerance (Paget et al. 2014; Pinto et al. 2025). The contribution of mitochondrial genome variation to this trait has also been suggested, based on analyses of interspecies hybrids (Baker et al. 2019; Li et al. 2019; Hewitt et al. 2020); exactly what the divergent mitotypes of *Saccharomyces* species do phenotypically, and how they exert their effects, remains incompletely understood.

We set out to pursue in more depth the phenotypic impact of mitochondrial variation between *Saccharomyces* species, both in isolation and in conjunction with nuclear loci. We anticipated that our findings would further illuminate the mechanisms of thermotolerance and their evolutionary history. We designed an approach that used *S. paradoxus*, a relatively thermosensitive species sister to *S. cerevisiae*, as a foreign background for manipulations of the nuclear and mitochondrial genomes, to investigate their functions and their interactions.

## Results

### *S. cerevisiae* mitotypes are sufficient to heighten respiratory behaviors in *S. paradoxus*

Previous studies have used yeast interspecies hybrids to establish an association between *S. cerevisiae* mitochondrial alleles and thermotolerance (Baker et al. 2019; Li et al. 2019; Hewitt et al. 2020), which parallels the advantage of *S. cerevisiae* purebreds at high temperature, relative to other species in the clade. We reasoned that investigating mitotype genetics in purebred *S. paradoxus* could help shed additional light on the mechanisms of evolutionary innovation along the *S. cerevisiae* lineage. For this purpose, we developed a series of cybrids, in which we introduced each mitochondrial genome in turn from a suite of *S. cerevisiae* isolates from different geographical locations, which all grew well in standard conditions and at high temperature (Figure S1), into a tester strain of *S. paradoxus*, the English oak tree isolate Z1 (Figure 1). Since they lack the endogenous *S. paradoxus* mitotype, these strains represented an experimental resource for the study of the phenotypic impact of *S. cerevisiae* mitochondrial alleles in an isogenic background, and they were viable and stable as expected (Sulo et al. 2003; Mário et al. 2015).

**Figure 1.**
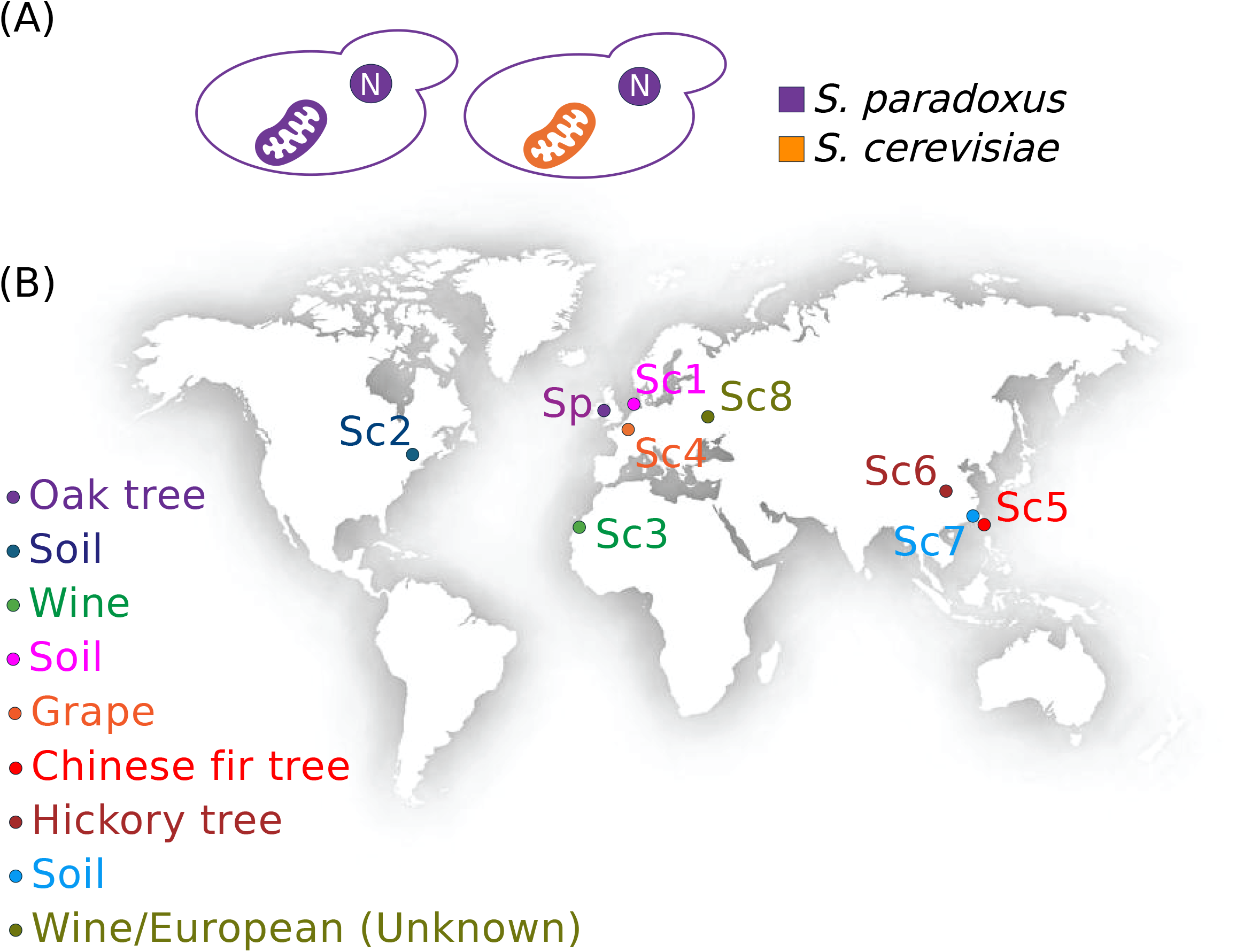
Experimental design and isolates for genetic dissection of *Saccharomyces* species variation in thermotolerance. Top, wild-type *S. paradoxus* (left) is used as the background for a cybrid (right) harboring the endogenous nuclear genome (N) and the mitochondrial genome from a *S. cerevisiae* donor (orange). Bottom, strains used in this work: Sp, *S. paradoxus* Z1; Sc1 to Sc8, *S. cerevisiae* DBVPG1373, YPS128, DBVPG6044, Y55, SJ6L01, SX3, GE14S01-7B, and DBVPG6765, respectively.

Before using our cybrids to address thermotolerance *per se*, we first investigated the hypothesis that respiration, as a proximal function of the mitochondrion and its genome, would be perturbed by species variation in mitotype. Wild-type *S. cerevisiae* strains as a rule grow better than those of *S. paradoxus* on glycerol, a non-fermentable carbon source (Figure S2), and in interspecies hybrid backgrounds, *S. cerevisiae* mitotypes promote growth in this condition (Li et al. 2019). To explore the impact of *S. cerevisiae* mitochondrial DNA alleles on respiratory growth in a purebred context, we assayed growth of our cybrids in liquid glycerol medium. The results revealed robust improvement attributable to *S. cerevisiae* mitotypes in the *S. paradoxus* background, for three of our *S. cerevisiae* donors (Figure 2A and Fig S3). The fourth cybrid, harboring mitochondrial DNA from the *S. cerevisiae* vineyard isolate Y55, had a defect in growth on glycerol (Figure 2A), which echoes previously reported incompatibilities of this mitotype in other backgrounds (Wolters et al. 2018). Together, these data suggested that pro-respiratory effects could be a prevalent characteristic of mitochondrial genomes from across the *S. cerevisiae* population.

**Figure 2.**
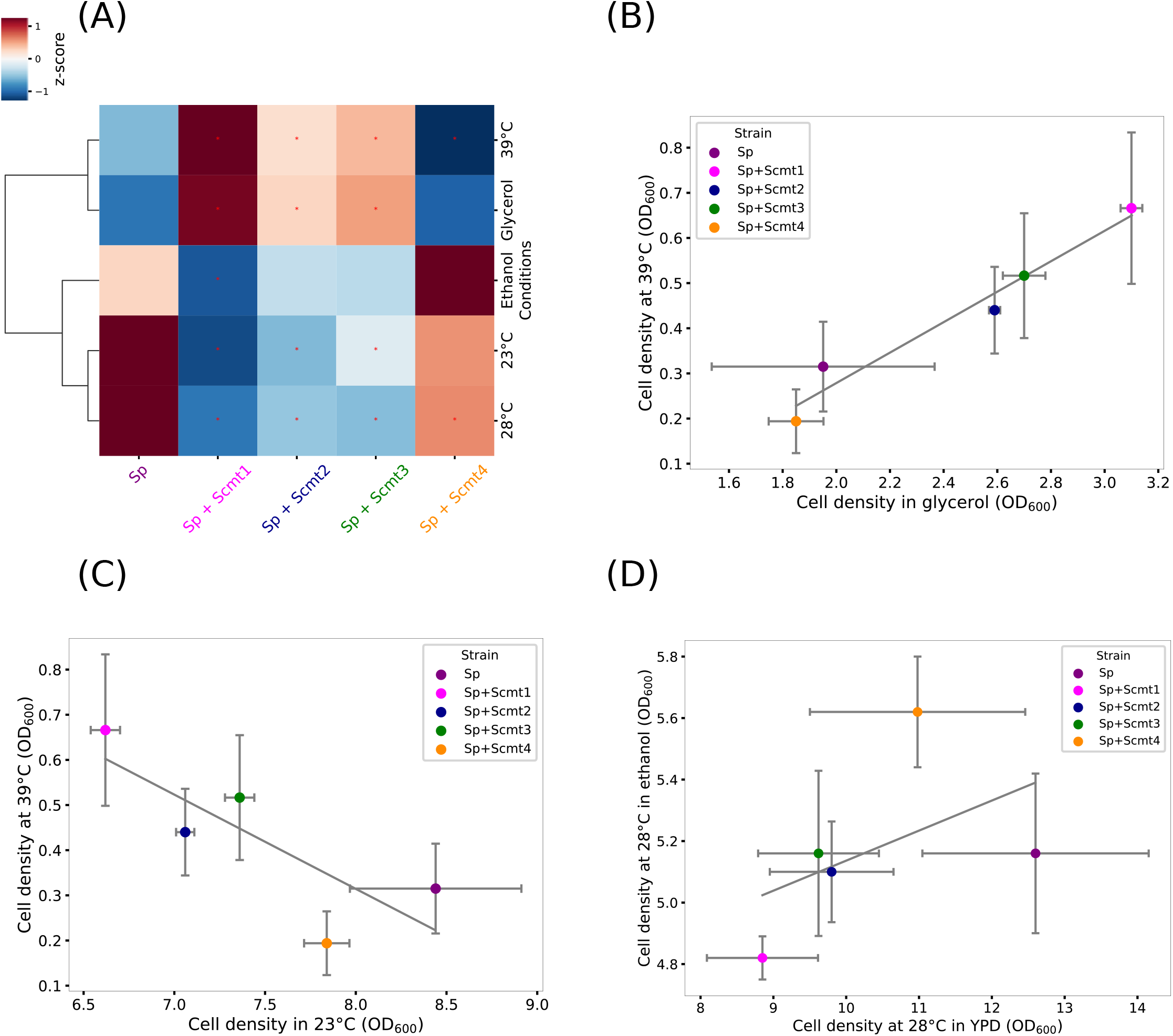
Condition-dependent growth and tradeoffs across cybrids. **(A)** Each cell reports cell density, as a mean across replicates (*n* ≥ 3), after 24 hours of growth in liquid culture in the respective condition, with glucose as the carbon source unless otherwise indicated. Each column reports results from wild-type *S. paradoxus* (Sp) or an isogenic cybrid harboring the mitochondrial genome from a *S. cerevisiae* donor (Scmt) labeled as in Figure 1. Colors report row-normalized z-score. **(B-D)** Shown are reanalyses of cell density measurements across strains from the cultures and conditions in **(A)**, with respective Spearman coefficients of −0.8, 1, and 0.67. In a given panel, each dot reports median across replicates (*n* ≥ 3), and error bars report standard deviation. Strain labels are as in **(A)**. *, Growth different from that of *S. paradoxus* at two-tailed Mann-Whitney *p* < 0.05. Raw growth measurements, Spearman correlation coefficients, and statistical analyses are reported in Table S2A to G.

As a more direct measure of metabolic effects driven by *S. cerevisiae* mitochondria DNA, we established an assay of reduction of the colorimetric substrate 1-(4,5-dimethylthiazol-2-yl)-3,5-diphenyltetrazolium bromide (MTT) in glucose medium. This signal serves as a reporter of activities of the electron transport chain, particularly succinate dehydrogenase (Slater et al. 1963; Sánchez and Königsberg 2006; Jain et al. 2018) (though see (Shoemaker et al. 2004) and (Peng et al. 2005)), and thus is informative as a marker of respiration. A first set of observations revealed no detectable impact of mitotype on MTT reduction during culture in liquid glucose medium at 28°C in the *S. paradoxus* background (Figure 3A and Fig S4A). We reasoned that the metabolic effect of *S. cerevisiae* mitotypes might be better resolved at higher temperatures, in which yeast cells respire particularly avidly (Rikhvanov et al. 2001). Consistent with this notion, in culture at 39°C in glucose medium, we observed increased MTT reduction in all cybrids except for that harboring *S. cerevisiae* Y55 mitochondrial DNA, relative to wild-type Z1 *S. paradoxus* with its endogenous mitochondrial genome (Figure 3B and Fig S4B). Together, these results established that in the *S. paradoxus* background, *S. cerevisiae* mitotypes can be sufficient for increased growth and metabolic behaviors that are signatures of respiration.

**Figure 3.**
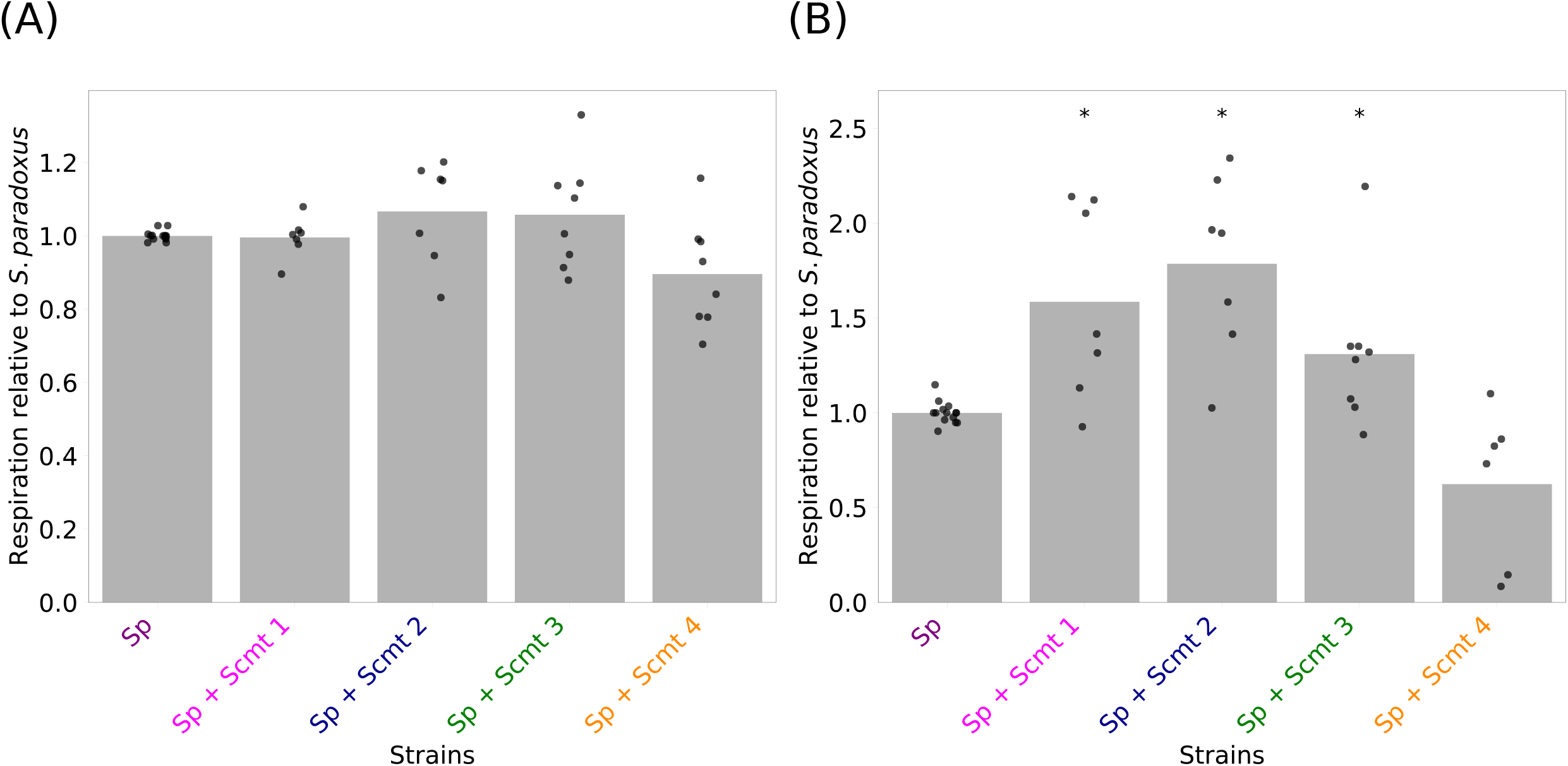
*S. cerevisiae* mitotypes boost respiration in the *S. paradoxus* nuclear background at 39°C. In each panel, in the main plot, the *y*-axis reports measurements of the reduction of 1-(4,5-dimethylthiazol-2-yl)-3,5-diphenyltetrazolium bromide after 24 hours of growth of a given strain in liquid culture with glucose as the carbon source at one temperature, normalized to the respective measurement wild-type *S. paradoxus*. In a given column, each point reports results from one biological replicate and the bar height reports the average. The *x*-axis denotes strain genotypes: wild-type *S. paradoxus* (Sp) or cybrids each harboring the mitochondrial genome from a *S. cerevisiae* donor (Scmt) labeled as in Figure 1. **(A)**, 28°C; **(B)**, 39°C. *, one-sample Wilcoxon *p* < 0.05. Raw data and statistical analyses are reported in Table S2H and I.

### *S. cerevisiae* mitotypes boost thermotolerance in *S. paradoxus*

We now turned to high-temperature growth proper, as an additional phenotype that we hypothesized would be modulated by mitotype in our yeast species comparison. To test the latter prediction, we measured cell density during growth in liquid glucose medium at 39°C, for each of our cybrid strains harboring *S. cerevisiae* mitochondrial DNA in Z1 *S. paradoxus*. Results revealed a 10% to 70% improvement in accumulated biomass in this assay, relative to wild-type *S. paradoxus*, in all cybrids except the one for which *S. cerevisiae* Y55 was the mitochondrial donor (Figure 2A). In an expanded panel of cybrids using additional *S. cerevisiae* mitotype donors, a boost in thermotolerance was also apparent in all cases (Figure S5). Thus, *S. cerevisiae* mitotypes were sufficient to confer thermotolerance benefits, in some cases sizeable ones, in *S. paradoxus*. Controls established that *S. cerevisiae* mitotypes had phenotypic effects in a fixed *S. cerevisiae* background mirroring those in *S. paradoxus*, though no *S. cerevisiae* cybrid exceeded the thermotolerance of the wild-type of this species (Figure S6). Interestingly, after 24 hours of incubation at 39°C, cultures of cybrids in the *S. paradoxus* background harbored very few viable cells (Figure S7). We conclude that in such cybrids, *S. cerevisiae* mitochondrial DNAs enable additional cell divisions during high-temperature exposure, after which failures of factors encoded by the *S. paradoxus* nuclear genome ultimately prevent further growth and kill cells outright.

Thermotolerance phenotypes as they varied across our panel of *S. paradoxus* cybrids correlated tightly with the respiratory behaviors we had documented in assays of growth in non-fermentable carbon sources in these strains (Figures 2A and 2B) and MTT reduction (Figure 3). That is, among *S. paradoxus* cybrids, those performing best in respiratory contexts were the most thermotolerant, and the worst in respiratory assays grew the worst at high temperature. These results are most consistent with a causal relationship between the traits, whereby the *S. cerevisiae* mitotypes that foster better thermotolerance do it by improving respiration—a metabolic underpinning for the capacity to grow under high-temperature challenge.

### Antagonistic pleiotropy by *S. cerevisiae* mitotypes in temperate conditions and ethanol

Previous studies using interspecies hybrids have reported a defect in cold temperatures attributable to *S. cerevisiae* mitotypes relative to those of *S. paradoxus* or other relatives (Baker et al. 2019; Li et al. 2019; Hewitt et al. 2020). To test whether such effects would be recapitulated in purebred backgrounds, in Z1 *S. paradoxus* derivatives carrying *S. cerevisiae* mitochondrial DNA, we assayed the carrying capacity after culture 23°C in liquid glucose medium, for comparison against a 28°C control and the high-temperature condition (39°C), also with glucose as a carbon source. The results revealed compromised cell density at 23°C in the cybrids that performed the best at high temperature (Figure 2A). This set up an almost perfect anticorrelation between 23°C growth and thermotolerance across strains (Figure 2C), mirroring the observation we had made with respiratory behaviors (Figures 2 and 3). We conclude that *S. cerevisiae* mitotypes have specialized to heat stress at the expense of performance at lower temperatures, suggestive of fine-tuning by evolution of the temperature optima of respiratory reactions encoded by the mitochondrial genome.

We also noted an anticorrelation across our cybrids between thermotolerance and cell density in glucose at 28°C: at the latter temperature, the most thermotolerant strains harboring *S. cerevisiae* DNA performed the worst (Figure 2A). To explore potential drivers of this pattern, we also assayed cell density at 28°C in medium with ethanol as the carbon source, in which we observed most cybrids carrying *S. cerevisiae* mitochondrial DNA to be at a disadvantage (Figure 2A). Growth patterns among strains at 28°C and in ethanol were robustly correlated (Figure 2D), suggestive of a mechanistic relationship between them, as expected given that *Saccharomyces* species catabolize ethanol late in the growth cycle after having secreted it (Piskur et al. 2006). In summary, based on our growth and respiration profiles, we conclude that *S. cerevisiae* mitochondrial genomes are drivers of a phenotypic syndrome that includes advantages in most facets of respiration as well as high-temperature growth, and disadvantages at cooler temperatures and during ethanol utilization.

### Positive epistasis between mitochondria and nuclear genomes in terms of thermotolerance

During the evolution of *S. cerevisiae* as a species, the pro-thermotolerance mitotypes we study here would have arisen alongside changes in nuclear genes which could modify their effects. We sought to shed light on these relationships by modeling them in the *S. paradoxus* background. To this end, we introduced *S. cerevisiae* mitochondrial genomes into a derivative of *S. paradoxus* Z1 that also harbored the alleles from the DBVPG1373 Dutch soil strain of *S. cerevisiae* at eight nuclear loci which contribute to thermotolerance (*AFG2, APC1, CEP3, DYN1, ESP1, MYO1, SCC2*, and *TAF2*, most of which encode cell division factors; (AlZaben et al. 2021)). In assays of cell density at 39°C, the combination of *S. cerevisiae* alleles in the nuclear and mitochondrial genomes together conferred a two-fold improvement relative to wild-type *S. paradoxus* (Figure 4, gray bars). This represented an increase in cell divisions early in the treatment, as no cybrids harboring *S. cerevisiae* nuclear alleles maintained viability after 24 hours at 39°C (Figure S8), paralleling results from cybrids in the wild-type *S. paradoxus* nuclear background (Figure S7). Advantages in cell density at 39°C manifested regardless of the origin of the mitotype: mitochondrial DNA from across our panel of *S. cerevisiae* strain donors yielded identical high-temperature growth in *S. paradoxus* in the presence of the nuclear thermotolerance genes from *S. cerevisiae*, including the Y55 mitotype that had compromised the phenotype in wild-type *S. paradoxus* (Figures 2A and 4). Thus, the latter incompatibility was resolved by *S. cerevisiae* alleles at our focal nuclear genes.

**Figure 4.**
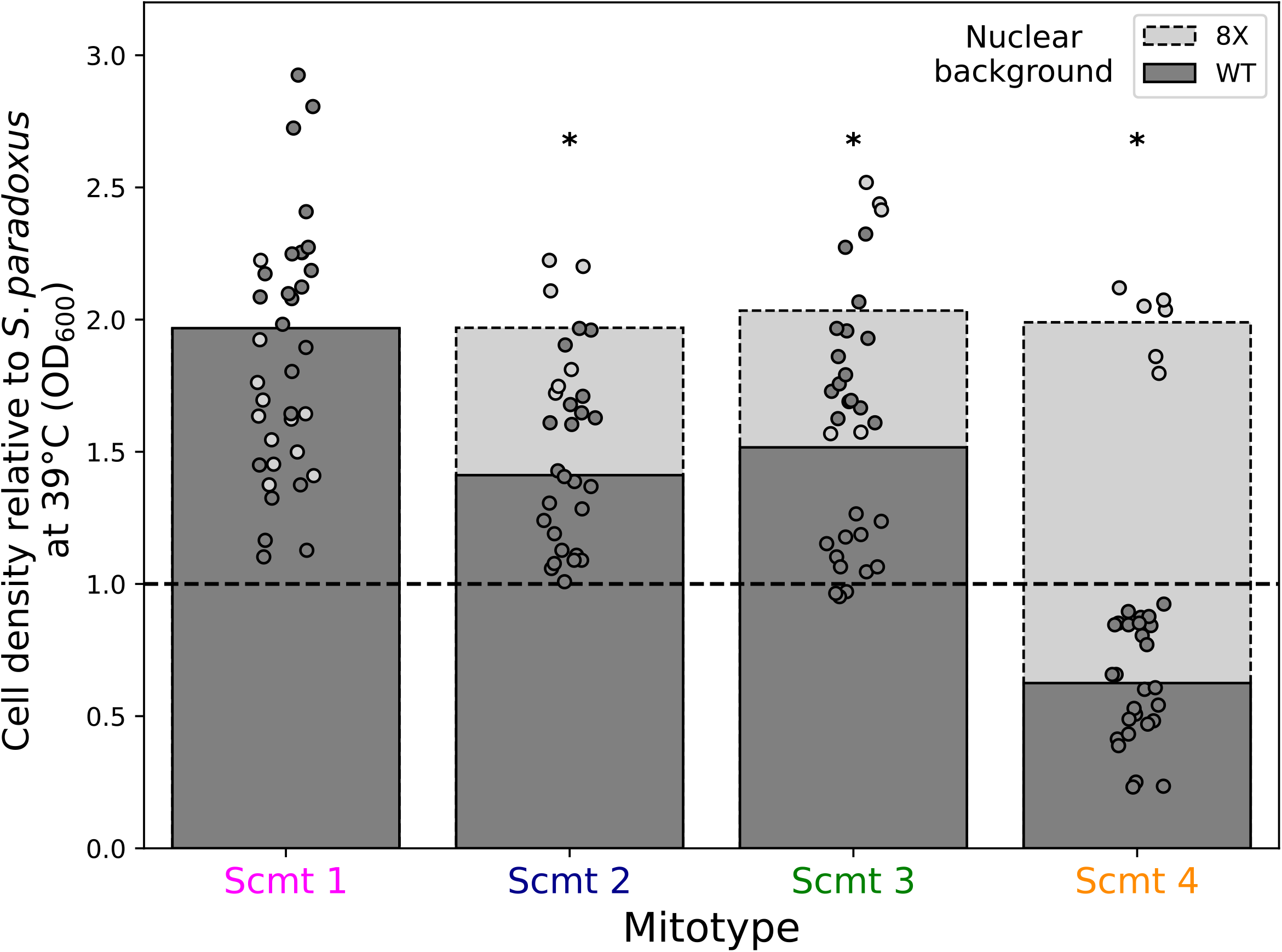
Epistasis between nuclear and mitochondrial *Saccharomyces* species variants that impact thermotolerance. In a given column, each color reports results from 24 hours of growth at 39°C, with glucose as the carbon source, of a strain of *S. paradoxus* with (8X) or without (WT) 8 thermotolerance loci from *S. cerevisiae*, harboring an *S. cerevisiae* mitotype. The *y*-axis reports the cell density of the indicated strain normalized to that of *S. paradoxus* with its native mitotype. Strain genotypes are as in Figure 1. Each point reports one replicate (*n* ≥ 6). *, the interaction term from a two-factor ANOVA with mitotype and nuclear genotype as factors was significant at *p* < 0.01. Raw growth measurements and statistical analyses are reported in Table S2A and J.

Also evident from our nuclear-mitochondrial transgenic strain panel was a pattern of positive epistasis. That is, the impact of each *S. cerevisiae* mitotype on thermotolerance was greater in the *S. paradoxus* derivative also harboring nuclear alleles from *S. cerevisiae* than in otherwise wild-type *S. paradoxus* (Figure 4; compare gray and black bars). Mitochondrial DNA from distinct *S. cerevisiae* donors, including Y55, all achieved peak performance at high temperature in the presence of *S. cerevisiae* nuclear genes; no such dependence was detectable in 28°C cultures (Figure S9). That said, the *S. cerevisiae* mitotype that promoted thermotolerance most strongly on its own in the *S. paradoxus* background, from DBVPG1373, benefited the least from the addition of DBVPG1373 alleles of nuclear genes (Figure 4, magenta). Relative to this transgenic, strains harboring other *S. cerevisiae* mitotypes, which had less impact on their own, achieved the same cell density at 39°C in the presence of DBVPG1373 nuclear alleles (Figure 4, blue, green, and orange). These results are most consistent with a ceiling beyond which any *S. cerevisiae* alleles of the mitochondrial genome, even when combined with *S. cerevisiae* alleles of our eight focal nuclear genes, are unable to improve thermotolerance, in the face of defects accruing from other *S. paradoxus* loci. Nonetheless, our data make clear that pro-thermotolerance alleles of mitochondrial DNAs and nuclear genes from *S. cerevisiae* together contribute to the phenotype in a manner that exceeds the sum of their parts.

## Discussion

A key goal of comparative genetics is to understand how evolution builds traits over long timescales. In this work, we have leveraged tractable *Saccharomyces* as a testbed for the study of interspecies trait genetics and evolution focused on the mitochondrial genome. Given the evidence that *S. cerevisiae* thermotolerance is the product of positive selection (Weiss et al. 2018; Abrams et al. 2021), it serves as a powerful model of the evolutionary genetics of adaptation over deep divergences. Our findings complement previous studies of mitotype effects in this system (Baker et al. 2019; Li et al. 2019; Hewitt et al. 2020) by tracing the phenotypes conferred by *S. cerevisiae* mitochondrial alleles in purebred backgrounds, including their genetic dependencies.

Our results shed new light on the genetic architecture and evolution of yeast thermotolerance. Our finding of a 70% boost in the trait by an *S. cerevisiae* mitotype outstrips the effect of any one of the nuclear loci previously reported in this system (Weiss et al. 2018). Sequence analyses revealed amino acid alleles unique to, and largely conserved in, *S. cerevisiae* in the *COX1* gene that represent candidate determinants of the mitochondrial effects we study here (Figure S11). But we have also seen that phenotypic impacts differ among *S. cerevisiae* mitotypes, consistent with the extensive precedent for polymorphism among mitochondrial genomes and their effects on human disease (Wallace 2015) as well as traits in other animal models (Mossman et al. 2016 Nov 14; Mossman et al. 2016; Patel et al. 2016; Tsai and St. John 2016; Camus et al. 2017; Camus and Dowling 2018; Mossman et al. 2019; Camus et al. 2020; Salminen and Vale 2020; Anderson et al. 2022; Brand et al. 2024; Serrano et al. 2024) and plants (Dubrovina and Kiselev 2016; Adhikari et al. 2019; Jo et al. 2019). Furthermore, none of our mitochondrial manipulations reconstituted more than a fraction of the phenotype of the *S. cerevisiae* wild-type in the *S. paradoxus* background, congruent with the very modest effects of *S. cerevisiae* nuclear alleles known to be adaptive in this system (Weiss et al. 2018; AlZaben et al. 2021). Thus, our findings deepen the emerging picture of a highly complex architecture for yeast thermotolerance, with mitochondrial variants and many nuclear loci of small to modest effect coming together along the *S. cerevisiae* lineage to build the adaptation.

Our data also make clear that alongside largely beneficial marginal effects, all but one of the *S. cerevisiae* mitotypes we study are even more advantageous at high temperature in the context of pro-thermotolerance nuclear alleles (the exception being the mitotype from Dutch soil isolate DBVPG1373). To date, such evidence for positive epistasis has been at a premium in the literature describing adaptive loci from the wild (Nelson and Grishin 2018; Syenina et al. 2020; Stern et al. 2022). This may be in part owing to a heavier emphasis in the field on negative epistasis in the evolutionarily difficult repacking of a given protein interior to achieve fitness (Weinreich et al. 2005; Starr and Thornton 2016), which could be more constrained than combinations of adaptive alleles of multiple proteins. And negative epistasis has been rife between mitochondrial and nuclear variants in analysis of traits that are not likely to promote fitness, including reproductive barriers and aging and disease (Ellison and Burton 2008; Arnqvist et al. 2010; Tranah 2011; Joseph et al. 2013; Lovell et al. 2015; Roux et al. 2016; Loewen and Ganetzky 2018; Wolters et al. 2018; Sujkowski et al. 2019; Nguyen et al. 2020; Bushel et al. 2022; Biot-Pelletier et al. 2023; Nguyen et al. 2023; Ibrahim et al. 2024). But even in the unlinked nuclear genes contributing to yeast thermotolerance, negative epistasis is the rule rather than the exception (AlZaben et al. 2021). As such, our data point to pro-thermotolerance mitoalleles as uniquely subject to positive epistasis with the nuclear genome in this adaptive system. Under one compelling model, these mitochondrial variants may have been especially easily accessible (Liao et al. 2022) and beneficial to populations during the evolution of *S. cerevisiae* thermotolerance, representing the so-called low-hanging fruit (Lenski 2017) of the adaptive process.

Though our data leave open the precise mechanism for such benefits, we propose that *S. cerevisiae* mitotypes produce extra energy and metabolites (Malecki et al. 2020) during thermal stress via enhanced respiration, relative to the putatively ancestral phenotypic program attributable to mitochondria from *S. paradoxus*. If so, an excess of basic building blocks would support the capacity of *S. cerevisiae* nuclear genes to confer their effects. The nuclear loci we study here govern housekeeping functions, namely cell division and transcription/translation, and some may prove to contribute more than others to the epistatic relationship with the mitochondrial genome.

Meanwhile, the growth defects we have seen by cybrids at lower temperatures dovetail with results from studies of *S. cerevisiae* nuclear thermotolerance loci and those using interspecies backgrounds (Baker et al. 2019; Li et al. 2019; Hewitt et al. 2020; AlZaben et al. 2021). As a general rule, evolutionary tradeoffs are routine in the adaptation literature, from prokaryotes (Abram et al. 2021; Backman et al. 2024; Dalldorf et al. 2024) to animals (Ou et al. 2020; Ackermans 2023; Metzler et al. 2023) and plants (Gao et al. 2022; He et al. 2022; Kurepa and Smalle 2023). For yeast in particular, the exact mechanism of antagonistic pleiotropy exhibited by thermotolerance loci in cooler conditions remains unclear. Exciting recent work has revealed that *S. cerevisiae* proteins, including those encoded by mitochondria, are more thermostable than those of sister species, establishing the likely biochemical underpinning of the trait (Walunjkar et al. 2025). But it may not follow that the *S. cerevisiae* alleles of the loci we study here compromise protein structure *per se* at low temperatures (Yang et al. 2015; Cui et al. 2024). Amino acid changes unique to *S. cerevisiae* could well have tuned the temperature optima for the function of many enzymes, as appears to be the case for the thermotolerance gene *DYN1* (Hong et al. 2016). Further work will be required to pinpoint exactly what biochemical mechanisms *S. cerevisiae* gave up as it acquired pro-thermotolerance alleles. As they stand, however, the genetics of this system help tell a story of a complex balancing act by evolution along the *S. cerevisiae* lineage, as nuclear and mitochondrial variants were assembled to convert a thermosensitive ancestor to a high-temperature specialist.

## Materials and methods

### Strain construction

Strains used in this study are listed in Table S1. For phenotyping in Figure S1, we used wild-type, homozygous diploid *S. cerevisiae* strains DBVPG1373, YPS128, DBVPG6044 and Y55, and *S. paradoxus* strains Z1, UFRJ50816, A4 and KPN3828, from the Saccharomyces Genome Resequencing Project (SGRP) collection (Table S1) (Cubillos et al. 2009). To make haploid strains for the recipients for *S. cerevisiae* mitochondrial DNA, we first generated prototrophic versions of diploid *S. paradoxus* Z1 and the *S. paradoxus* harboring the eight *S. cerevisiae* loci, designated as 8X (AlZaben et al. 2021). For this, the *HO* loci of the diploid wild-type Z1 and 8X strains were knocked out using the hygromycin resistance gene as the selection marker using the transformation protocol from (Reuß et al. 2004), with selection on 200 μg/ml hygromycin and PCR and sequence confirmation. The resulting *hoΔ*::Hyg diploid strains were induced to sporulate essentially as in (Lee et al. 2008), and mating types of the resulting haploids were checked via halo assay following the protocol in (Hoffman et al. 2002), using JRY02375 and JRY02176 as the mating testers. Next, for haploid MATα *S. paradoxus* Z1 and its descendant 8X, we generated a petite version of each (lacking mitochondrial DNA, ρ^0^), using the protocol adopted from (Amine et al. 2021). In short, yeast cells were streaked from −80°C stocks on yeast peptone dextrose (YPD) plates and grown at 28°C for 2 days. A colony was inoculated into 3 ml of YPD and incubated at 28°C overnight. 10^4^ cells were inoculated into 1 ml of YPD supplemented with ethidium bromide (EtBr, 10 μg/ml) and incubated at 28°C overnight. Cells were harvested and washed twice with ddH_2_O. About 200 cells were plated onto YPD plates and grown at 28°C for 2-3 days to form visible colonies. The plates were replicated to yeast peptone glycerol (YPG) plates to look for colonies that lost the ability to respire, *i*.*e*. those in the ρ^0^ state.

Finally, we used wild-type, MATα haploid, G418-resistant, hygromycin-resistant *S. cerevisiae* strains DBVPG1373, YPS128, DBVPG6044, Y55 and DBVPG6765 from the SGRP collection (Cubillos et al. 2009), and MATα haploid hygromycin-resistant *S. cerevisiae* strains (SJ6L01, Sx3 and GE14S01-7B), a generous gift from Joseph Schacherer (Table S1) (Peter et al. 2018), as donors of mitochondrial DNA for cybrids in *S. paradoxus* wild-type (Z1), 8X (Z1 with 8 *S. cerevisiae* nuclear loci), and *S. cerevisiae* (DBVPG1373) backgrounds. Our method for cybrid construction used the cytoduction protocol of (Nguyen et al. 2020) as follows. We first constructed a heterokaryon-competent *S. cerevisiae* strain harboring each mitotype of interest, referred to as a carrier. For this, a single colony of each donor in turn, and a single colony of the ρ^0^ *kar1*-d15 mutant (JRY5450, a generous gift from Dr. Jasper Rine), were each inoculated into a separate 3 mL of liquid YPD (1% yeast extract, 2% peptone, 2% glucose) and incubated at 28°C overnight with shaking at 250 rpm. We mixed 100 μL of the two, spotted on a YPD plate, and incubated at 28°C for 4 hours. The mixed cellular patch was inoculated into 3 mL of YPD and incubated at 28°C for 2 hours with shaking at 250 rpm to promote cell division. After a 2-hour incubation, 10^-5^ OD_600_ of cells were spread on a YPG plate (1% yeast extract, 2% peptone, 2% glycerol) to select for mitochondrial genome retention. Colonies on the YPG plates were transferred to a YPD plate with 200 μg/mL hygromycin or 200 μg/mL G418 to check for hygromycin and G418 sensitivity; we earmarked and preserved drug-sensitive, respiration-competent strains as those in which the donor mitochondrial DNA had successfully been transferred to the JRY5450 nuclear background. We next used each carrier strain in turn to transfer the *S. cerevisiae* mitochondrial DNA of interest into the haploid *S. paradoxus* Z1 ρ^0^ strain and, separately, its descendant, the 8X ρ^0^ strain. This cytoduction proceeded as above except that the carrier and the *S. paradoxus* strain were mated and the progeny were selected on YPG alone; afterward we subjected them to the halo assay, to eliminate the carrier-*S. paradoxus* diploid hybrids, and only those strains that behaved as haploid MATα were retained as cybrids.

### Growth assays

Measurements of cell density were done as described (AlZaben et al. 2021) with some modifications. To make sure the cells were respiratory-competent before the assay, strains were streaked on YPG plates and were allowed to grow at 28°C for 4 days to form colonies. Three colonies were inoculated into 15 mL of liquid YPG and incubated at 28°C for overnight with shaking at 250 rpm. The cultures reached 0.4-0.8 OD_600_ after a 16-hour growth; in these pre-cultures, assays of glycogen and trehalose storage (see below) detected no difference between genotypes (Figure S10). The cultures were then back-diluted into 5 mL of YPD, YPE (1% yeast extract, 2% peptone, 2% ethanol), or YPG to achieve 0.1 OD_600_/mL, and then incubated at 23°C, 28°C or 39°C for 24 hours. This procedure, from streaking on solid agar plates to biomass measurement, was repeated at least twice for each strain with three technical repeats in each biological replicate. We collected the readouts of cell density from all the replicates across all days for a given strain and compared each mitochondrial transgenic to *S. paradoxus* with a two-tailed Mann-Whitney U test. For epistasis analysis, a two-way ANOVA test was carried out to examine the impact of the genotype-mitotype interaction on thermotolerance. All statistical analyses were done with scipy.stats in Python 3.7.

### MTT assay

To measure respiration efficiency, pre-culture and treatment were as in Growth assays, above, and after 24 hours we carried out the MTT assay as in (Sánchez and Königsberg 2006). Independent experiments from growth to MTT measurement were performed at least six times. The readouts were normalized to that of wild-type *S. paradoxus* at either 28°C or 39°C. Two-tailed one-sample Wilcoxon test was performed using the normalized data with a hypothesized value of 1 with scipy.stats in Python 3.7.

### Growth of *S. cerevisiae* and *S. paradoxus* in glycerol

Data are from (Liti et al. 2009) and Mann-Whitney U test was performed to compare the growths of the two species in glycerol with scipy.stats in Python 3.7.

### Trehalose and glycogen assays

To measure the contents of trehalose and glycogen stored in yeast, we took advantage of the protocol from (Chen and Futcher 2017). Briefly, yeast cells were pre-grown in YPG overnight. 0.3 OD_600_ of cells were harvested, and the pellets were washed with 1 ml of ddH_2_O to remove residual media. The pellets were resuspended in 125 μl of 0.25 M Na_2_CO_3_ solution, and incubated at 95°C for 3 hours in PCR tubes. 75 μl of 1 M acetic acid and 300 μl of 0.2 M sodium acetate, pH 5.2 were added to the mix. The mix was immediately divided into two 250-μl aliquots. For glycogen measurements, 10 μl of *Aspergillus niger* α-amyloglucosidase (20 mg/ml in 0.2 M sodium acetate, pH 5.2) was added to one of the aliquots, and incubated at 57°C overnight. For trehalose measurements, 15 μl of 0.2 M sodium acetate, pH 8 was added to the other aliquot to adjust the pH to ~5.8, and then 3 μl of porcine trehalase was added to the sample, and incubated at 37°C overnight. After overnight incubation, the content of glucose, digested from either trehalose or glycogen, in each sample was measured using Glucose (GO) Assay Kit (Cat. No. GAGO20) at absorbance of 540nm. OD_540_ readouts in this setting represent the content of glucose in each sample, reporting the concentration of the substrates, trehalose and glycogen, are in the cells.

### Multiple sequence alignment

*S. cerevisiae* mitochondrial genomes were from (De Chiara et al. 2020). North American *S. paradoxus* mitochondrial genomes (NCBI accessions: KY287641-KY287662) were from (Leducq et al. 2017), and European and Asian *S. paradoxus* genomes (NCBI accessions: JQ862335.1, KP712799.1, KP712802.1, KP712786.1, KP712796.1) were from (Procházka et al. 2012). Multiple sequence alignment was performed using Muscle v. 5.1. (Edgar 2022), the allele frequency was calculated and plotted with Python 3.7, and the alignment was visualized with AliView v. 3.0 (Larsson 2014).

## Supplementary figure captions

**Figure S1. Stark growth differences between *S. cerevisiae* and *S. paradoxus* at high temperature**. In a given panel, the *y*-axis reports cell density after 24 hours of growth in liquid culture of a wild-type homozygous diploid *Saccharomyces* strain in one temperature condition, with glucose as the carbon source. Points reports technical replicates (*n* = 3) and bar heights report means. Sc1, Sc2, Sc3 and Sc4 denote *S. cerevisiae* DBVPG1373, YPS128, DBVPG6044 and Y55, respectively. Sp1, Sp2, Sp3 and Sp4 denote *S. paradoxus* KPN3828, UFRJ50816, A4 and Z1, respectively. **(A)** 28°C; **(B)** 39°C. *, two-tailed Mann-Whitney *p* < 0.05. Raw growth measurements and statistical analyses are reported in Table S2K.

**Figure S2. Wild-type strain and species variation in growth on glycerol**. In a given panel, each bar reports results from the indicated wild-type *S. cerevisiae* or *S. paradoxus* strain grown in liquid culture with glycerol as the carbon source, normalized with respect to a standard reference-strain control, from (Warringer et al. 2011). The *y*-axis reports relative growth rate **(A)** and efficiency **(B)**. *, two-tailed Mann-Whitney *p* < 0.05. Raw growth measurements and statistical analyses are reported in Table S2L to S2N.

**Figure S3. Condition-dependent growth across *Saccharomyces* species and cybrids**. Data are as in Figure 2A except that wild-type *S. cerevisiae* (DBVPG1373) is included. Raw growth measurements and statistical analyses are reported in Table S2A to S2G.

**Figure S4. *S. cerevisiae* mitotypes boost respiration in the *S. paradoxus* nuclear background at 39°C with addition of *S. cerevisiae***. Data are as in Figure 3 except that wild-type *S. cerevisiae* (DBVPG1373) is included. Raw MTT measurements and statistical analyses are reported in Table S2H and S2I.

**Figure S5. *S. cerevisiae* mitotypes are sufficient to boost thermotolerance in *S. paradoxus***. Each column reports results from *S. paradoxus* harboring mitochondrial DNA from a *S. cerevisiae* strain. The *y*-axis reports cell density after 24 hours of growth at 39°C of a cybrid normalized to that of the wild-type *S. paradoxus*, which is shown as the dashed line. *, two-tailed Mann-Whitney *p* < 0.05. Bar height reports average across replicates, and each dot is one technical or biological replicate. Scmt 1 to Scmt 8 denote mitotypes from Sc 1 to Sc 8 in Figure 1. Raw growth measurements and statistical analyses are reported in Table S2A and S2O.

**Figure S6. Effects of *S. cerevisiae* mitotypes in *S. cerevisiae* on thermotolerance echo those in *S. paradoxus***. Each column reports results from *S. cerevisiae* harboring mitochondrial DNA from a *S. cerevisiae* strain. The *y*-axis reports cell density after 24 hours of growth at 39°C. *, two-tailed Mann-Whitney *p* < 0.05. Bar height reports average across replicates, and each dot is one technical or biological replicate. Labels on the *x*-axis are as in Figure 2A. Raw growth measurements and statistical analyses are reported in Table S2A and S2J.

**Figure S7. Chronic 39°C treatment is lethal to *S. paradoxus* and its cybrids**. Shown are results of plating of the indicated strains after 24 hours of culture in one temperature condition, with glucose as the carbon source, followed by incubation of plates at 28°C for two days. Strains were wild-type *S. cerevisiae* (Sc), wild-type *S. paradoxus* (Sp), or a cybrid in the *S. paradoxus* background harboring the mitochondrial genome from a *S. cerevisiae* donor (Scmt) labeled as in Figure 1 of the main text.

**Figure S8. Chronic 39°C treatment is lethal to *S. paradoxus* harboring *S. cerevisiae* nuclear thermotolerance loci and to its cybrids**. Labels are as in Figure S7 except that the strains were wild-type *S. cerevisiae* (Sc), wild-type *S. paradoxus* (Sp), *S. paradoxus* harboring eight nuclear loci from *S. cerevisiae* (8X), or a cybrid in the 8X background harboring the mitochondrial genome from a *S. cerevisiae* donor (Scmt) labeled as in Figure 1 of the main text.

**Figure S9. No detectable epistasis between nuclear and mitochondrial *Saccharomyces* species variants that impact 28°C growth**. Data are as Figure 4 of the main text except that cell densities were measured after 24 hours of growth at 28°C. Raw growth measurements and statistical analyses are reported in Table S2A and S2J.

**Figure S10. No detectable differences in glycogen and trehalose storage in *S. paradoxus* cybrids**. In a given panel, each bar reports energy storage by cells of the indicated strain after growth with glycerol as the carbon source at 28°C, either glycogen (A) or trehalose (B). Each dot denotes one biological or technical replicate. Labels on the *x*-axis are as in Figure 2A of the main text. Raw trehalose and glycogen measurements and statistical analyses are reported in Table S2P and S2Q.

**Figure S11. A multiple sequence alignment analysis shows highly diverged amino acid variants in *COX1* between *S. paradoxus* and *S. cerevisiae***. At top, each line reports one *COX1* amino acid sequence of *S. cerevisiae* (737 strains) and *S. paradoxus* (27 strains) as denoted on the left. At bottom, bar heights in each panel report the frequency of the indicated amino acid allele in the indicated species. Not shown are infrequent alleles in *S. cerevisia*e., namely N (0.3%) and T (0.1%) at position 55 and premature stop codons (0.3%) at position 58. Mitochondrial genome accessions are cited in the Materials and Methods section.

## Supplementary table captions

**Table S1. Strains used in this work**.

**Table S2. Growth data and statistics**. D1373, YPS128, D6044 and Y55 indicate *S. cerevisiae* strains DBVPG1373, YPS128, DBVPG6044 and Y55, respectively; Sp, *S. paradoxus* Z1. Table S2A contains all OD_600_ measurements of each strain at 39°C in standard glucose medium. Table S2B reports the statistical analysis of the OD_600_ measurements of the cybrids in **(A)** compared to wild-type *S. paradoxus* or the *S. paradoxus* harboring *S. cerevisiae* nuclear loci (8X) by Mann-Whitney U test. Table S2C contains all OD_600_ measurements of each strain at 28°C in standard glucose medium. Table S2D contains all OD_600_ measurements of each strain at 28°C in medium with glycerol as a carbon source. Table S2E contains all OD_600_ measurements of each strain at 28°C in medium with ethanol as a carbon source. Table S2F contains all OD_600_ measurements of each strain at 23°C in standard glucose medium. Table S2G reports the Spearman correlation coefficients and respective *p*-values of Figures 2B, 2C and 2D. Table 2H reports the raw readouts of the MTT assay at 28°C and 39°C. Table 2I reports the *p*-values of the normalized MTT measurements by one-sample Wilcoxon with a hypothesized value of 1. Tabel S2J reports the *p*-values of two-way ANOVA of the impact of the interaction between *S. cerevisiae* mitotypes and the 8 *S. cerevisiae* loci in the *S. paradoxus* background on high temperature growth (Figure 4). Table S2K reports reports the OD_600_ measurements of wild-type *S. cerevisiae* and *S. paradoxus* strains at 28°C and 39°C in standard glucose medium with the statistical analysis (Figure S1). Table S2L and S2M report the growth efficiency and rate in glycerol as the carbon source from (Warringer et al. 2011). Table S2N reports the *p*-values of glycerol growth efficiency and rate of the wild-type *S. cerevisiae* and *S. paradoxus* strains by two-tailed Mann-Whitney U test (Warringer et al. 2011) (Figure S2). Table S2O reports the *p*-values of Figure S5 by two-tailed Mann-Whitney U test. Table S2P and S2Q report the raw readouts of trehalose and glycogen assays and *p*-values by two-tailed Mann-Whitney U test, respectively (Figure S10).

## References

Abram F, Arcari T, Guerreiro D, O’Byrne CP. 2021. Evolutionary trade-offs between growth and survival: The delicate balance between reproductive success and longevity in bacteria. In: Advances in Microbial Physiology. Vol. 79. Elsevier; p 133–162 https://linkinghub.elsevier.com/retrieve/pii/S006529112100014X. 10.1016/bs.ampbs.2021.07.002

Abrams MB et al. 2021. Population and comparative genetics of thermotolerance divergence between yeast species Gresham D, editor. G3. 11(7):jkab139. 10.1093/g3journal/jkab139

Ackermans NL. 2023. Neurobiological tradeoffs of headbutting bovids. Trends in Neurosciences. 46(11):898–900. 10.1016/j.tins.2023.08.004

Adhikari B, Caruso CM, Case AL. 2019. Beyond balancing selection: frequent mitochondrial recombination contributes to high-female frequencies in gynodioecious Lobelia siphilitica (Campanulaceae). New Phytologist. 224(3):1381–1393. 10.1111/nph.16136

Allen Orr H. 2001. The genetics of species differences. Trends in Ecology & Evolution. 16(7):343–350. 10.1016/S0169-5347(01)02167-X

AlZaben F, Chuong JN, Abrams MB, Brem RB. 2021. Joint effects of genes underlying a temperature specialization tradeoff in yeast Fay JC, editor. PLoS Genet. 17(9):e1009793. 10.1371/journal.pgen.1009793

Amine AAA et al. 2021. Experimental evolution improves mitochondrial genome quality control in Saccharomyces cerevisiae and extends its replicative lifespan. Current Biology. 31(16):3663-3670.e4. 10.1016/j.cub.2021.06.026

Anderson L et al. 2022. Variation in mitochondrial DNA affects locomotor activity and sleep in Drosophila melanogaster. Heredity. 129(4):225–232. 10.1038/s41437-022-00554-w

Arnqvist G et al. 2010. Genetic architecture of metabolic rate: environment specific epistasis between mitochondrial and nuclear genes in an insect. Evolution. 64(12):3354–3363. 10.1111/j.1558-5646.2010.01135.x

Backman T, Burbano HA, Karasov TL. 2024. Tradeoffs and constraints on the evolution of tailocins. Trends in Microbiology. 32(11):1084–1095. 10.1016/j.tim.2024.04.001

Baker EP et al. 2019. Mitochondrial DNA and temperature tolerance in lager yeasts. Sci Adv. 5(1):eaav1869. 10.1126/sciadv.aav1869

Bazzicalupo A. 2022. Local adaptation in fungi. FEMS Microbiology Reviews. 46(6):fuac026. 10.1093/femsre/fuac026

Biot-Pelletier D et al. 2023. Evolutionary Trajectories are Contingent on Mitonuclear Interactions Zhang J, editor. Molecular Biology and Evolution. 40(4):msad061. 10.1093/molbev/msad061

Bozdag GO et al. 2021. Breaking a species barrier by enabling hybrid recombination. Current Biology. 31(4):R180–R181. 10.1016/j.cub.2020.12.038

Brand JA, Garcia-Gonzalez F, Dowling DK, Wong BBM. 2024. Mitochondrial genetic variation as a potential mediator of intraspecific behavioural diversity. Trends in Ecology & Evolution. 39(2):199–212. 10.1016/j.tree.2023.09.009

Bushel PR et al. 2022. Mitochondrial-nuclear epistasis underlying phenotypic variation in breast cancer pathology. Sci Rep. 12(1):1393. 10.1038/s41598-022-05148-4

Camus MF, Dowling DK. 2018. Mitochondrial genetic effects on reproductive success: signatures of positive intrasexual, but negative intersexual pleiotropy. Proc Biol Sci. 285(1879):20180187. 10.1098/rspb.2018.0187

Camus MF, O’Leary M, Reuter M, Lane N. 2020. Impact of mitonuclear interactions on life-history responses to diet. Phil Trans R Soc B. 375(1790):20190416. 10.1098/rstb.2019.0416

Camus MF, Wolff JN, Sgrò CM, Dowling DK. 2017. Experimental Support That Natural Selection Has Shaped the Latitudinal Distribution of Mitochondrial Haplotypes in Australian Drosophila melanogaster. Mol Biol Evol. 34(10):2600–2612. 10.1093/molbev/msx184

Cao Y et al. 2022. Sex differences in heart mitochondria regulate diastolic dysfunction. Nat Commun. 13(1):3850. 10.1038/s41467-022-31544-5

Chen Y, Futcher B. 2017. Assaying Glycogen and Trehalose in Yeast. Bio Protoc. 7(13):e2371. 10.21769/BioProtoc.2371

Chou J-Y et al. 2010. Multiple Molecular Mechanisms Cause Reproductive Isolation between Three Yeast Species Noor MAF, editor. PLoS Biol. 8(7):e1000432. 10.1371/journal.pbio.1000432

Chou J-Y, Leu J-Y. 2015. The Red Queen in mitochondria: cyto-nuclear co-evolution, hybrid breakdown and human disease. Front Genet. 6 [accessed 2025 Feb 18]. http://www.frontiersin.org/Evolutionary_and_Population_Genetics/10.3389/fgene.2015.00187/abstract. 10.3389/fgene.2015.00187

Crandall JG, Fisher KJ, Sato TK, Hittinger CT. 2023. Ploidy evolution in a wild yeast is linked to an interaction between cell type and metabolism Zanders SE, editor. PLoS Biol. 21(11):e3001909. 10.1371/journal.pbio.3001909

Cubillos FA, Louis EJ, Liti G. 2009. Generation of a large set of genetically tractable haploid and diploid Saccharomyces strains. FEMS Yeast Research. 9(8):1217–1225. 10.1111/j.1567-1364.2009.00583.x

Cui S, Furukawa R, Akanuma S. 2024. Insights into the low-temperature adaptation of an enzyme as studied through ancestral sequence reconstruction. [accessed 2024 Dec 16]. http://biorxiv.org/lookup/doi/10.1101/2024.09.05.611558. 10.1101/2024.09.05.611558

Dalldorf C et al. 2024. The hallmarks of a tradeoff in transcriptomes that balances stress and growth functions Fodor A, editor. mSystems. 9(7):e00305–24. 10.1128/msystems.00305-24

De Chiara M et al. 2020. Discordant evolution of mitochondrial and nuclear yeast genomes at population level. BMC Biol. 18(1):49. 10.1186/s12915-020-00786-4

Dubrovina AS, Kiselev KV. 2016. Age-associated alterations in the somatic mutation and DNA methylation levels in plants Weber A, editor. Plant Biol J. 18(2):185–196. 10.1111/plb.12375

Edgar RC. 2022. Muscle5: High-accuracy alignment ensembles enable unbiased assessments of sequence homology and phylogeny. Nat Commun. 13(1):6968. 10.1038/s41467-022-34630-w

Elena SF. 2017. Local adaptation of plant viruses: lessons from experimental evolution. Molecular Ecology. 26(7):1711–1719. 10.1111/mec.13836

Ellison CK, Burton RS. 2008. Genotype-dependent variation of mitochondrial transcriptional profiles in interpopulation hybrids. Proc Natl Acad Sci USA. 105(41):15831–15836. 10.1073/pnas.0804253105

Ferreira T, Rodriguez S. 2024. Mitochondrial DNA: Inherent Complexities Relevant to Genetic Analyses. Genes. 15(5):617. 10.3390/genes15050617

Forejt J, Jansa P. 2023. Meiotic Recognition of Evolutionarily Diverged Homologs: Chromosomal Hybrid Sterility Revisited Malik H, editor. Molecular Biology and Evolution. 40(4):msad083. 10.1093/molbev/msad083

Fredericks LR et al. 2021. The Species-Specific Acquisition and Diversification of a K1-like Family of Killer Toxins in Budding Yeasts of the Saccharomycotina. PLoS Genet. 17(2):e1009341. 10.1371/journal.pgen.1009341

Gao H et al. 2022. Natural variations of ZmSRO1d modulate the trade-off between drought resistance and yield by affecting ZmRBOHC-mediated stomatal ROS production in maize. Molecular Plant. 15(10):1558–1574. 10.1016/j.molp.2022.08.009

Gonçalves P et al. 2011. Evidence for Divergent Evolution of Growth Temperature Preference in Sympatric Saccharomyces Species Chaturvedi V, editor. PLoS ONE. 6(6):e20739. 10.1371/journal.pone.0020739

He Z, Webster S, He SY. 2022. Growth–defense trade-offs in plants. Current Biology. 32(12):R634–R639. 10.1016/j.cub.2022.04.070

Hénault M, Marsit S, Charron G, Landry CR. 2022. Hybridization drives mitochondrial DNA degeneration and metabolic shift in a species with biparental mitochondrial inheritance. Genome Res. 32(11–12):2043–2056. 10.1101/gr.276885.122

Hewitt SK et al. 2020. Plasticity of Mitochondrial DNA Inheritance and its Impact on Nuclear Gene Transcription in Yeast Hybrids. Microorganisms. 8(4):494. 10.3390/microorganisms8040494

Hoffman GA, Garrison TR, Dohlman HG. 2002. Analysis of RGS Proteins in Saccharomyces cerevisiae. In: Methods in Enzymology. Vol. 344. Elsevier; p 617–631 [accessed 2024 Dec 17]. https://linkinghub.elsevier.com/retrieve/pii/S0076687902447441. 10.1016/S0076-6879(02)44744-1

Hong W et al. 2016. The Effect of Temperature on Microtubule-Based Transport by Cytoplasmic Dynein and Kinesin-1 Motors. Biophysical Journal. 111(6):1287–1294. 10.1016/j.bpj.2016.08.006

Hou J, Friedrich A, de Montigny J, Schacherer J. 2014. Chromosomal Rearrangements as a Major Mechanism in the Onset of Reproductive Isolation in Saccharomyces cerevisiae. Current Biology. 24(10):1153–1159. 10.1016/j.cub.2014.03.063

Houtkooper RH et al. 2013. Mitonuclear protein imbalance as a conserved longevity mechanism. Nature. 497(7450):451–457. 10.1038/nature12188

Ibrahim R, Bahilo Martinez M, Dobson AJ. 2024. Rapamycin’s lifespan effect is modulated by mito-nuclear epistasis in Drosophila. Aging Cell. e14328. 10.1111/acel.14328

Jain AK et al. 2018. Models and Methods for In Vitro Toxicity. In: In Vitro Toxicology. Elsevier; p 45–65 [accessed 2025 Feb 25]. https://linkinghub.elsevier.com/retrieve/pii/B9780128046678000031. 10.1016/B978-0-12-804667-8.00003-1

Jhuang H, Lee H, Leu J. 2017. Mitochondrial–nuclear co-evolution leads to hybrid incompatibility through pentatricopeptide repeat proteins. EMBO Reports. 18(1):87–101. 10.15252/embr.201643311

Jo YD et al. 2019. Mitotypes Based on Structural Variation of Mitochondrial Genomes Imply Relationships With Morphological Phenotypes and Cytoplasmic Male Sterility in Peppers. Front Plant Sci. 10:1343. 10.3389/fpls.2019.01343

Joseph B et al. 2013. Cytoplasmic genetic variation and extensive cytonuclear interactions influence natural variation in the metabolome. eLife. 2:e00776. 10.7554/eLife.00776

Kellis M et al. 2003. Sequencing and comparison of yeast species to identify genes and regulatory elements. Nature. 423(6937):241–254. 10.1038/nature01644

Kraemer SA, Boynton PJ. 2017. Evidence for microbial local adaptation in nature. Molecular Ecology. 26(7):1860–1876. 10.1111/mec.13958

Kurepa J, Smalle JA. 2023. Plant Hormone Modularity and the Survival-Reproduction Trade-Off. Biology. 12(8):1143. 10.3390/biology12081143

Larsson A. 2014. AliView: a fast and lightweight alignment viewer and editor for large datasets. Bioinformatics. 30(22):3276–3278. 10.1093/bioinformatics/btu531

Leducq J-B et al. 2017. Mitochondrial Recombination and Introgression during Speciation by Hybridization. Molecular Biology and Evolution. 34(8):1947–1959. 10.1093/molbev/msx139

Lee H-Y et al. 2008. Incompatibility of Nuclear and Mitochondrial Genomes Causes Hybrid Sterility between Two Yeast Species. Cell. 135(6):1065–1073. 10.1016/j.cell.2008.10.047

Lenski RE. 2017. Experimental evolution and the dynamics of adaptation and genome evolution in microbial populations. The ISME Journal. 11(10):2181–2194. 10.1038/ismej.2017.69

Li XC et al. 2019. Mitochondria-encoded genes contribute to evolution of heat and cold tolerance in yeast. Sci Adv. 5(1):eaav1848. 10.1126/sciadv.aav1848

Liao S, Chen L, Song Z, He H. 2022. The fate of damaged mitochondrial DNA in the cell. Biochimica et Biophysica Acta (BBA) - Molecular Cell Research. 1869(5):119233. 10.1016/j.bbamcr.2022.119233

Liti G et al. 2009. Population genomics of domestic and wild yeasts. Nature. 458(7236):337– 341. 10.1038/nature07743

Loewen CA, Ganetzky B. 2018. Mito-Nuclear Interactions Affecting Lifespan and Neurodegeneration in a Drosophila Model of Leigh Syndrome. Genetics. 208(4):1535–1552. 10.1534/genetics.118.300818

Lovell JT et al. 2015. Exploiting Differential Gene Expression and Epistasis to Discover Candidate Genes for Drought-Associated QTLs in Arabidopsis thaliana. Plant Cell. 27(4):969– 983. 10.1105/tpc.15.00122

Lupo O, Krieger G, Jonas F, Barkai N. 2021. Accumulation of cis- and trans-regulatory variations is associated with phenotypic divergence of a complex trait between yeast species Andrews B, editor. G3 Genes|Genomes|Genetics. 11(2):jkab016. 10.1093/g3journal/jkab016

Malecki M, Kamrad S, Ralser M, Bähler J. 2020. Mitochondrial respiration is required to provide amino acids during fermentative proliferation of fission yeast. EMBO Reports. 21(11):e50845. 10.15252/embr.202050845

Mário Š, Poláková S, Jatzová K, Sulo P. 2015. Post-zygotic sterility and cytonuclear compatibility limits in S. cerevisiae xenomitochondrial cybrids. Front Genet. 5 [accessed 2024 Nov 7]. http://journal.frontiersin.org/article/10.3389/fgene.2014.00454/abstract. 10.3389/fgene.2014.00454

Marsit S et al. 2021. The neutral rate of whole-genome duplication varies among yeast species and their hybrids. Nat Commun. 12(1):3126. 10.1038/s41467-021-23231-8

Melvin RG, Ballard JWO. 2006. Intraspecific variation in survival and mitochondrial oxidative phosphorylation in wild-caught Drosophila simulans. Aging Cell. 5(3):225–233. 10.1111/j.1474-9726.2006.00211.x

Meng X et al. 2022. GWAS on multiple traits identifies mitochondrial ACONITASE3 as important for acclimation to submergence stress. Plant Physiology. 188(4):2039–2058. 10.1093/plphys/kiac011

Metzler S, Kirchner J, Grasse AV, Cremer S. 2023. Trade-offs between immunity and competitive ability in fighting ant males. BMC Ecol Evo. 23(1):37. 10.1186/s12862-023-02137-7

Moran BM et al. 2024. A lethal mitonuclear incompatibility in complex I of natural hybrids. Nature. 626(7997):119–127. 10.1038/s41586-023-06895-8

Mossman JA et al. 2016. Mitonuclear Interactions Mediate Transcriptional Responses to Hypoxia in Drosophila. Mol Biol Evol. msw246. 10.1093/molbev/msw246

Mossman JA, Biancani LM, Rand DM. 2019. Mitochondrial genomic variation drives differential nuclear gene expression in discrete regions of Drosophila gene and protein interaction networks. BMC Genomics. 20(1):691. 10.1186/s12864-019-6061-y

Mossman JA, Biancani LM, Zhu C-T, Rand DM. 2016. Mitonuclear Epistasis for Development Time and Its Modification by Diet in Drosophila. Genetics. 203(1):463–484. 10.1534/genetics.116.187286

Nelson ED, Grishin NV. 2018. Inference of epistatic effects in a key mitochondrial protein. Phys Rev E. 97(6):062404. 10.1103/PhysRevE.97.062404

Nguyen THM et al. 2020. Mitochondrial-nuclear coadaptation revealed through mtDNA replacements in Saccharomyces cerevisiae. BMC Evol Biol. 20(1):128. 10.1186/s12862-020-01685-6

Nguyen THM et al. 2023. Mapping mitonuclear epistasis using a novel recombinant yeast population Hopper AK, editor. PLoS Genet. 19(3):e1010401. 10.1371/journal.pgen.1010401

Nikolov LA, Tsiantis M. 2015. Interspecies Gene Transfer as a Method for Understanding the Genetic Basis for Evolutionary Change: Progress, Pitfalls, and Prospects. Front Plant Sci. 6:1135. 10.3389/fpls.2015.01135

Oppong RF et al. 2022. Personality traits are consistently associated with blood mitochondrial DNA copy number estimated from genome sequences in two genetic cohort studies. eLife. 11:e77806. 10.7554/eLife.77806

Ou Q et al. 2020. Evolutionary trade-off in reproduction of Cambrian arthropods. Sci Adv. 6(18):eaaz3376. 10.1126/sciadv.aaz3376

Paget CM, Schwartz J-M, Delneri D. 2014. Environmental systems biology of cold-tolerant phenotype in Saccharomyces species adapted to grow at different temperatures. Mol Ecol. 23(21):5241–5257. 10.1111/mec.12930

Pakendorf B, Stoneking M. 2005. MITOCHONDRIAL DNA AND HUMAN EVOLUTION. Annu Rev Genom Hum Genet. 6(1):165–183. 10.1146/annurev.genom.6.080604.162249

Parra M et al. 2023. Assembly and comparative genome analysis of a Patagonian Aureobasidium pullulans isolate reveals unexpected intraspecific variation. Yeast. 40(5–6):197– 213. 10.1002/yea.3853

Patel MR et al. 2016. A mitochondrial DNA hypomorph of cytochrome oxidase specifically impairs male fertility in Drosophila melanogaster. eLife. 5:e16923. 10.7554/eLife.16923

Peng L, Wang B, Ren P. 2005. Reduction of MTT by flavonoids in the absence of cells. Colloids and Surfaces B: Biointerfaces. 45(2):108–111. 10.1016/j.colsurfb.2005.07.014

Peris D et al. 2023. Macroevolutionary diversity of traits and genomes in the model yeast genus Saccharomyces. Nat Commun. 14(1):690. 10.1038/s41467-023-36139-2

Peter J et al. 2018. Genome evolution across 1,011 Saccharomyces cerevisiae isolates. Nature. 556(7701):339–344. 10.1038/s41586-018-0030-5

Pinto J, Balarezo-Cisneros LN, Delneri D. 2025. Exploring adaptation routes to cold temperatures in the Saccharomyces genus Peris D, editor. PLoS Genet. 21(2):e1011199. 10.1371/journal.pgen.1011199

Piskur J et al. 2006. How did Saccharomyces evolve to become a good brewer? Trends in Genetics. 22(4):183–186. 10.1016/j.tig.2006.02.002

Procházka E, Franko F, Poláková S, Sulo P. 2012. A complete sequence of Saccharomyces paradoxus mitochondrial genome that restores the respiration in S. cerevisiae. FEMS Yeast Res. 12(7):819–830. 10.1111/j.1567-1364.2012.00833.x

Quéméneur J-B et al. 2022. The relationships between growth rate and mitochondrial metabolism varies over time. Sci Rep. 12(1):16066. 10.1038/s41598-022-20428-9

Rees JS, Castellano S, Andrés AM. 2020. The Genomics of Human Local Adaptation. Trends in Genetics. 36(6):415–428. 10.1016/j.tig.2020.03.006

Reuß O, Vik Å, Kolter R, Morschhäuser J. 2004. The SAT1 flipper, an optimized tool for gene disruption in Candida albicans. Gene. 341:119–127. 10.1016/j.gene.2004.06.021

Rikhvanov EG et al. 2001. [No title found]. Microbiology. 70(4):462–465. 10.1023/A:1010442429489

Roop JI, Chang KC, Brem RB. 2016. Polygenic evolution of a sugar specialization trade-off in yeast. Nature. 530(7590):336–339. 10.1038/nature16938

Roux F et al. 2016. Cytonuclear interactions affect adaptive traits of the annual plant Arabidopsis thaliana in the field. Proc Natl Acad Sci USA. 113(13):3687–3692. 10.1073/pnas.1520687113

Salminen TS, Vale PF. 2020. Drosophila as a Model System to Investigate the Effects of Mitochondrial Variation on Innate Immunity. Front Immunol. 11:521. 10.3389/fimmu.2020.00521

Salvadó Z et al. 2011. Temperature Adaptation Markedly Determines Evolution within the Genus Saccharomyces. Appl Environ Microbiol. 77(7):2292–2302. 10.1128/AEM.01861-10

Sánchez NS, Königsberg M. 2006. Using yeast to easily determine mitochondrial functionality with 1-(4,5-dimethylthiazol-2-yl)-3,5-diphenyltetrazolium bromide (MTT) assay. Biochem Molecular Bio Educ. 34(3):209–212. 10.1002/bmb.2006.49403403209

Serrano IM et al. 2024. Mitochondrial haplotype and mito-nuclear matching drive somatic mutation and selection throughout ageing. Nat Ecol Evol. 8(5):1021–1034. 10.1038/s41559-024-02338-3

Shoemaker M, Cohen I, Campbell M. 2004. Reduction of MTT by aqueous herbal extracts in the absence of cells. Journal of Ethnopharmacology. 93(2–3):381–384. 10.1016/j.jep.2004.04.011

Slater TF, Sawyer B, Sträuli U. 1963. Studies on succinate-tetrazolium reductase systems. Biochimica et Biophysica Acta. 77:383–393. 10.1016/0006-3002(63)90513-4

Smukowski Heil C et al. 2021. Transposable Element Mobilization in Interspecific Yeast Hybrids Vieira C, editor. Genome Biology and Evolution. 13(3):evab033. 10.1093/gbe/evab033

Sork VL. 2017. Genomic Studies of Local Adaptation in Natural Plant Populations. Journal of Heredity. 109(1):3–15. 10.1093/jhered/esx091

Starr TN, Thornton JW. 2016. Epistasis in protein evolution. Protein Science. 25(7):1204–1218. 10.1002/pro.2897

Stern DB, Anderson NW, Diaz JA, Lee CE. 2022. Genome-wide signatures of synergistic epistasis during parallel adaptation in a Baltic Sea copepod. Nat Commun. 13(1):4024. 10.1038/s41467-022-31622-8

Sujkowski A et al. 2019. Mito-nuclear interactions modify Drosophila exercise performance. Mitochondrion. 47:188–205. 10.1016/j.mito.2018.11.005

Sullivan LB, Chandel NS. 2014. Mitochondrial reactive oxygen species and cancer. Cancer Metab. 2(1):17. 10.1186/2049-3002-2-17

Sulo P et al. 2003. The efficiency of functional mitochondrial replacement in species has directional character. FEMS Yeast Research. 4(1):97–104. 10.1016/S1567-1356(03)00109-0

Sun J et al. 2019. Mitochondrial variation in small brown planthoppers linked to multiple traits and probably reflecting a complex evolutionary trajectory. Molecular Ecology. 28(14):3306– 3323. 10.1111/mec.15148

Swamy KBS et al. 2022. Proteotoxicity caused by perturbed protein complexes underlies hybrid incompatibility in yeast. Nat Commun. 13(1):4394. 10.1038/s41467-022-32107-4

Sweeney J, Kuehne H, Sniegowski P. 2004. Sympatric natural and populations have different thermal growth profiles. FEMS Yeast Research. 4(4–5):521–525. 10.1016/S1567-1356(03)00171-5

Syenina A et al. 2020. Positive epistasis between viral polymerase and the 3′ untranslated region of its genome reveals the epidemiologic fitness of dengue virus. Proc Natl Acad Sci USA. 117(20):11038–11047. 10.1073/pnas.1919287117

Tranah GJ. 2011. Mitochondrial–nuclear epistasis: Implications for human aging and longevity. Ageing Research Reviews. 10(2):238–252. 10.1016/j.arr.2010.06.003

Tsai T, St. John JC. 2016. The role of mitochondrial DNA copy number, variants, and haplotypes in farm animal developmental outcome. Domestic Animal Endocrinology. 56:S133–S146. 10.1016/j.domaniend.2016.03.005

Vijayraghavan S et al. 2019. Mitochondrial Genome Variation Affects Multiple Respiration and Nonrespiration Phenotypes in Saccharomyces cerevisiae. Genetics. 211(2):773–786. 10.1534/genetics.118.301546

Wallace DC. 2015. Mitochondrial DNA Variation in Human Radiation and Disease. Cell. 163(1):33–38. 10.1016/j.cell.2015.08.067

Walunjkar N et al. 2025. Pervasive Divergence in Protein Thermostability is Mediated by Both Structural Changes and Cellular Environments Zhang J, editor. Molecular Biology and Evolution. 42(7):msaf137. 10.1093/molbev/msaf137

Warringer J et al. 2011. Trait Variation in Yeast Is Defined by Population History Kruglyak L, editor. PLoS Genet. 7(6):e1002111. 10.1371/journal.pgen.1002111

Weinreich DM, Watson RA, Chao L. 2005. Perspective: Sign epistasis and genetic constraint on evolutionary trajectories. Evolution. 59(6):1165–1174

Weiss CV et al. 2018. Genetic dissection of interspecific differences in yeast thermotolerance. Nat Genet. 50(11):1501–1504. 10.1038/s41588-018-0243-4

Weiss CV, Brem RB. 2019. Dissecting Trait Variation across Species Barriers. Trends in Ecology & Evolution. 34(12):1131–1136. 10.1016/j.tree.2019.07.013

Wolff JN, Ladoukakis ED, Enríquez JA, Dowling DK. 2014. Mitonuclear interactions: evolutionary consequences over multiple biological scales. Philos Trans R Soc Lond B Biol Sci. 369(1646):20130443. 10.1098/rstb.2013.0443

Wolters JF et al. 2018. Mitochondrial Recombination Reveals Mito–Mito Epistasis in Yeast. Genetics. 209(1):307–319. 10.1534/genetics.117.300660

Wolters JF, Chiu K, Fiumera HL. 2015. Population structure of mitochondrial genomes in Saccharomyces cerevisiae. BMC Genomics. 16(1):451. 10.1186/s12864-015-1664-4

Yang L-L, Tang S-K, Huang Y, Zhi X-Y. 2015. Low Temperature Adaptation Is Not the Opposite Process of High Temperature Adaptation in Terms of Changes in Amino Acid Composition. Genome Biol Evol. 7(12):3426–3433. 10.1093/gbe/evv232

