## Supplementary figures and images for "The role of mitotype variation and positive epistasis in trait differences between *Saccharomyces* species"

### Supplemental figure 1

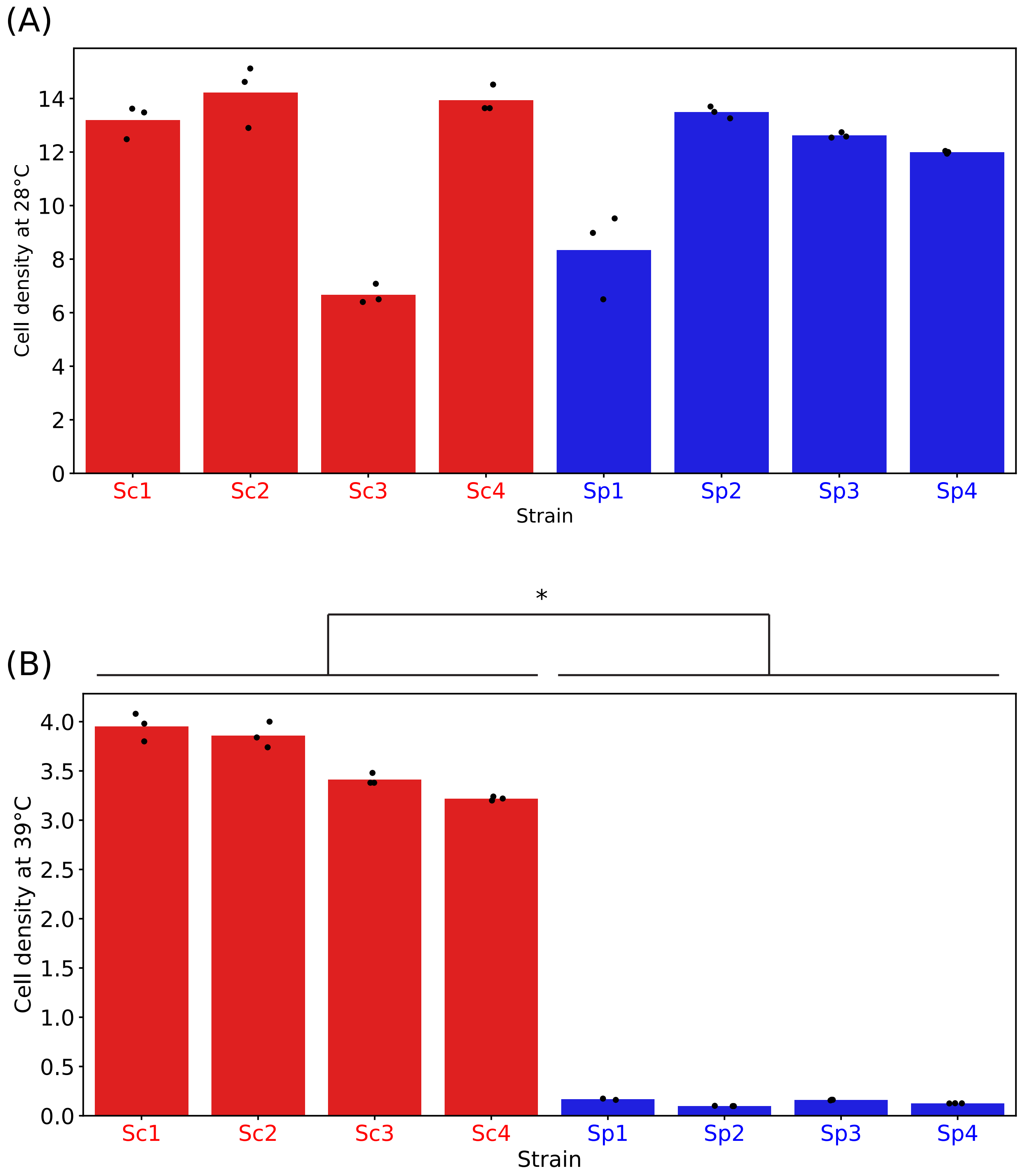

### Supplemental figure 2

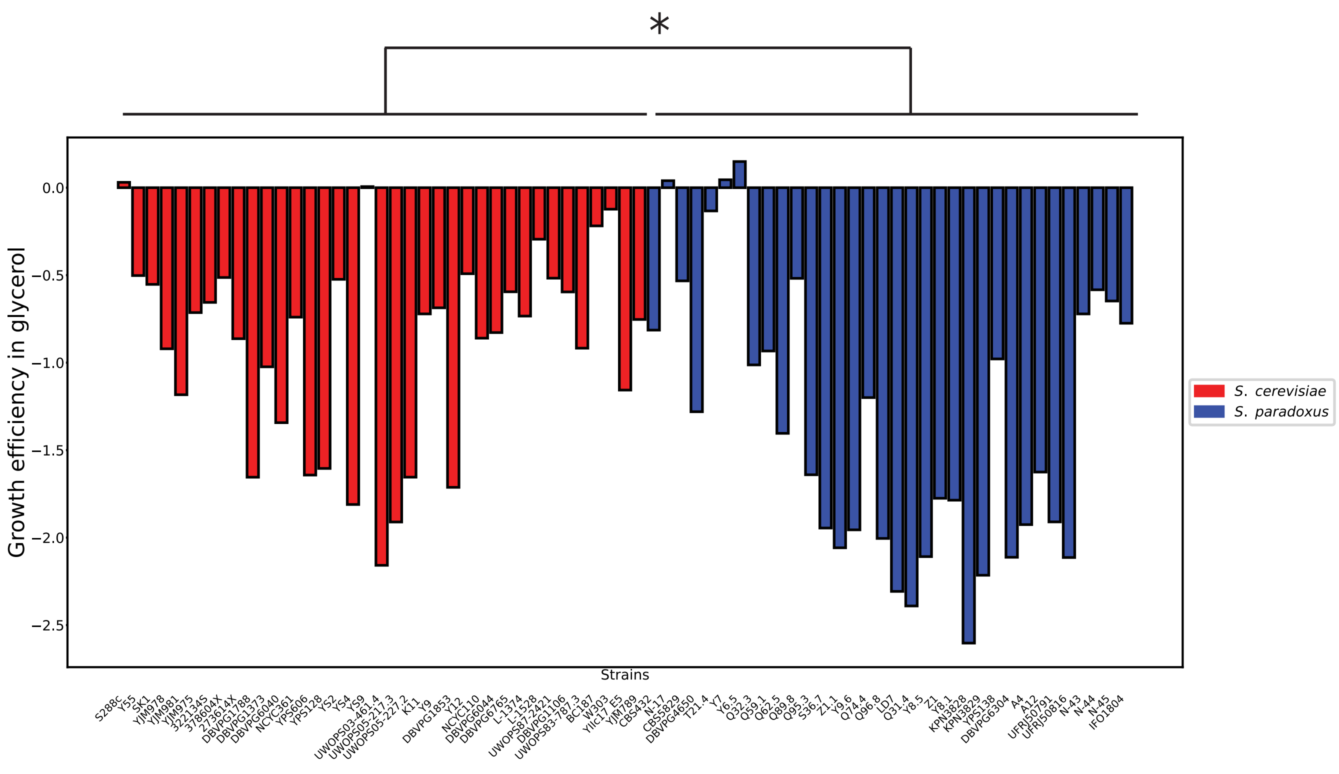

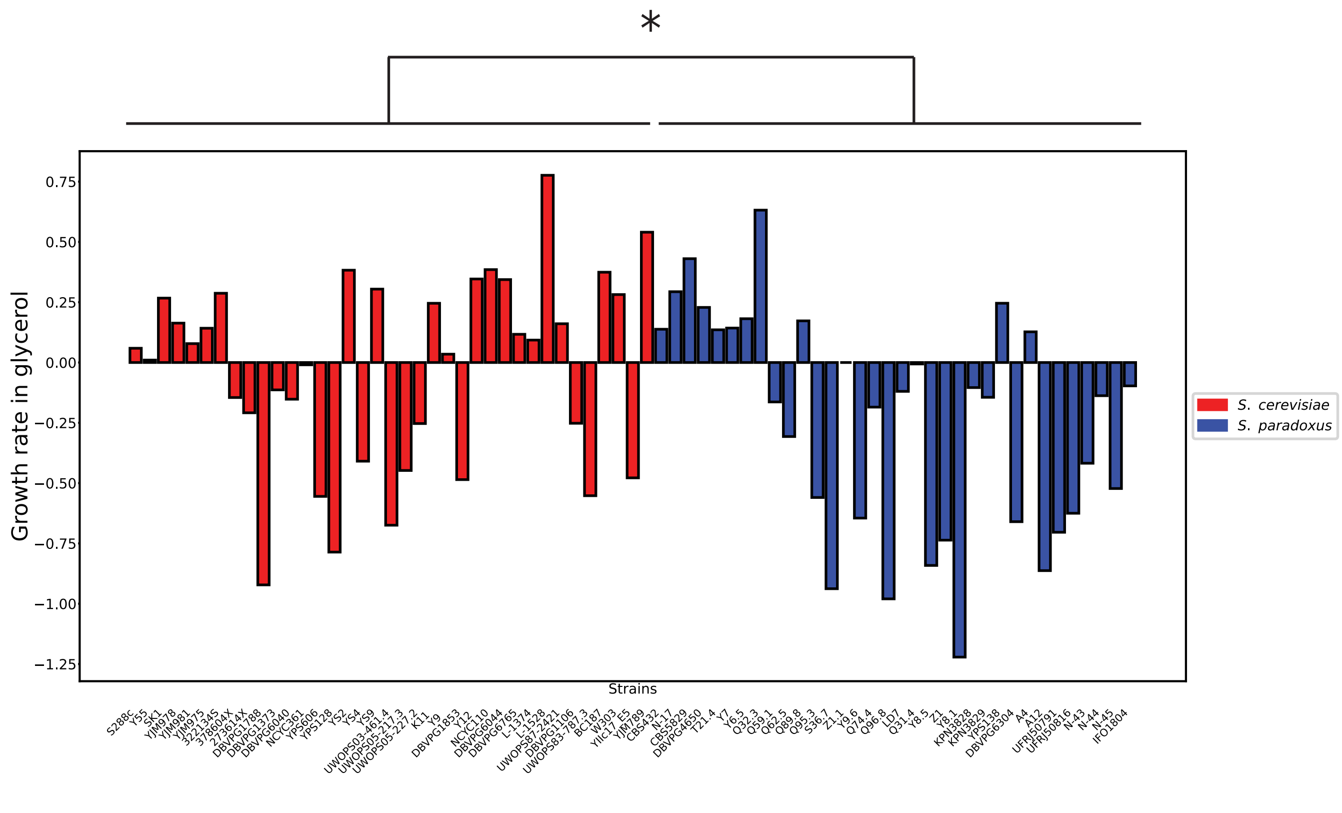

### Supplemental figure 3

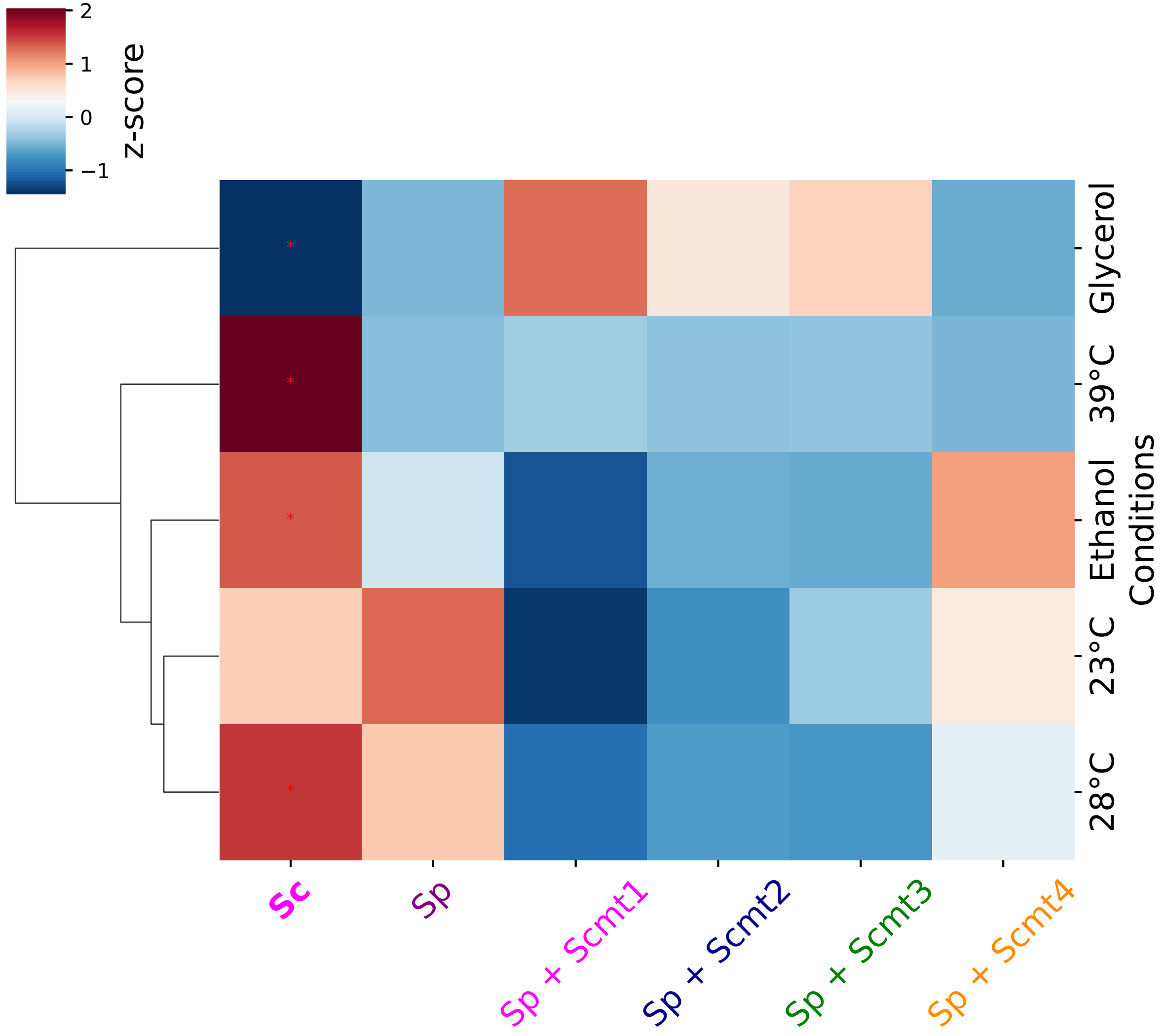

### Supplemental figure 4

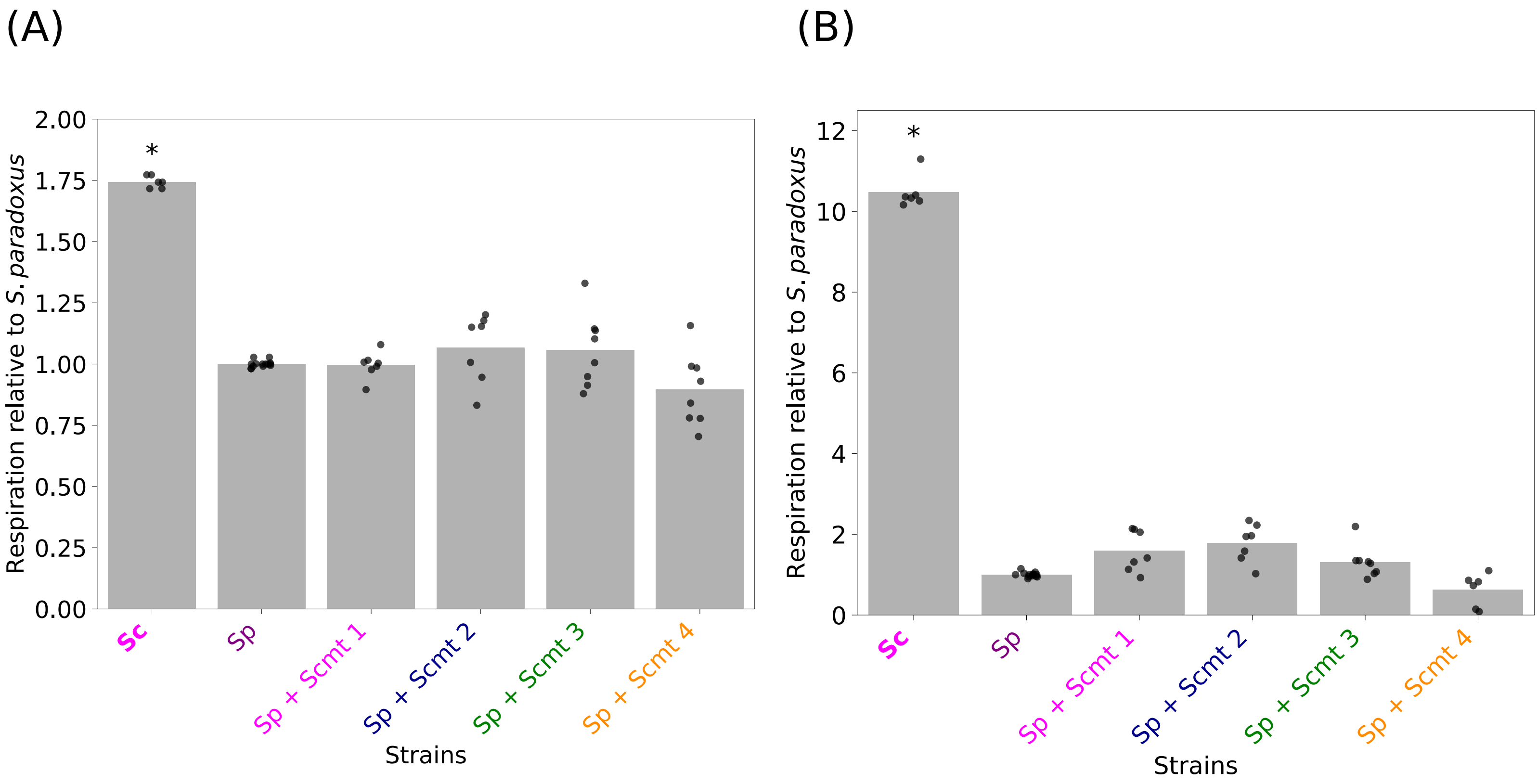

### Supplemental figure 5

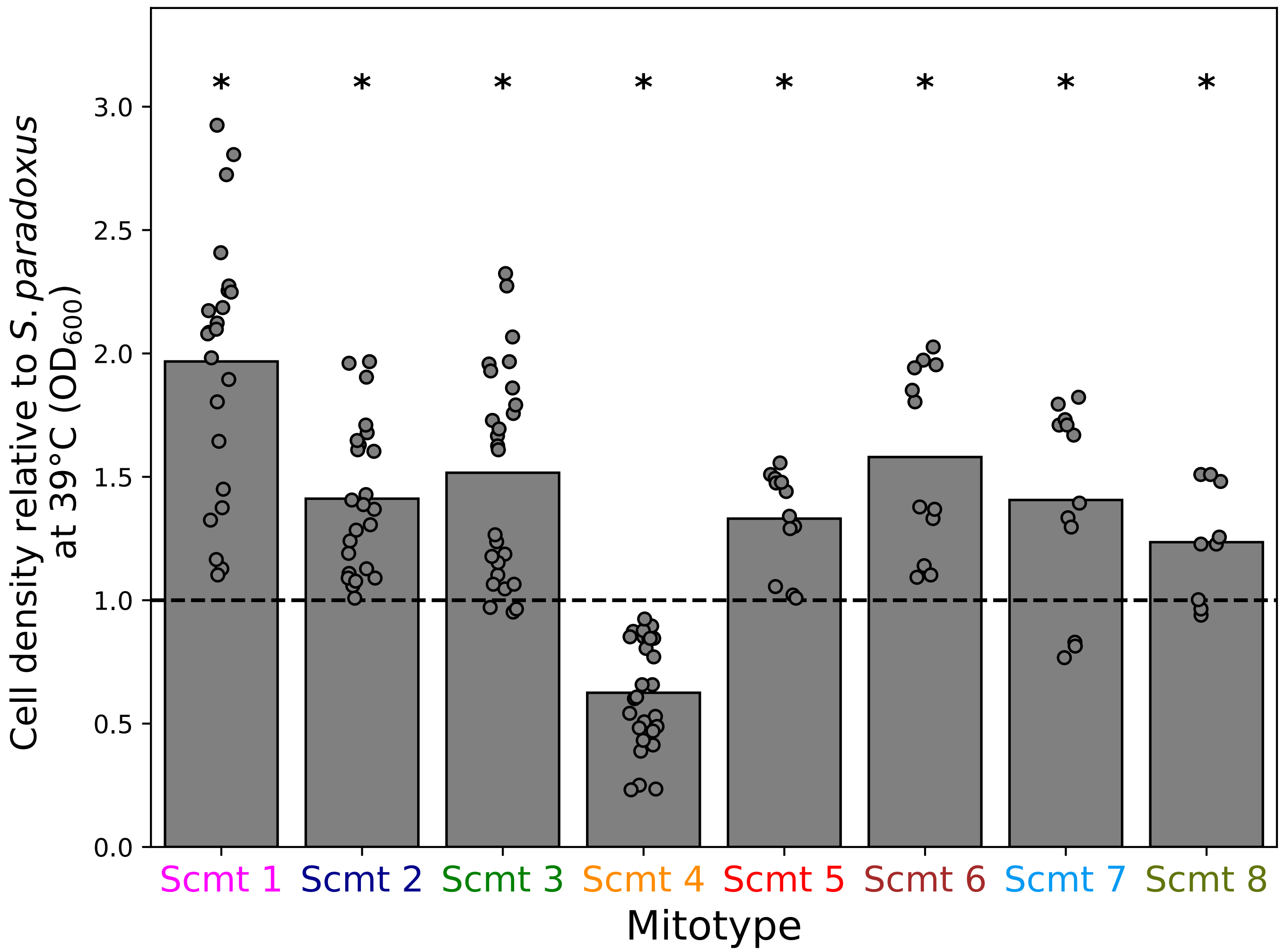

### Supplemental figure 6

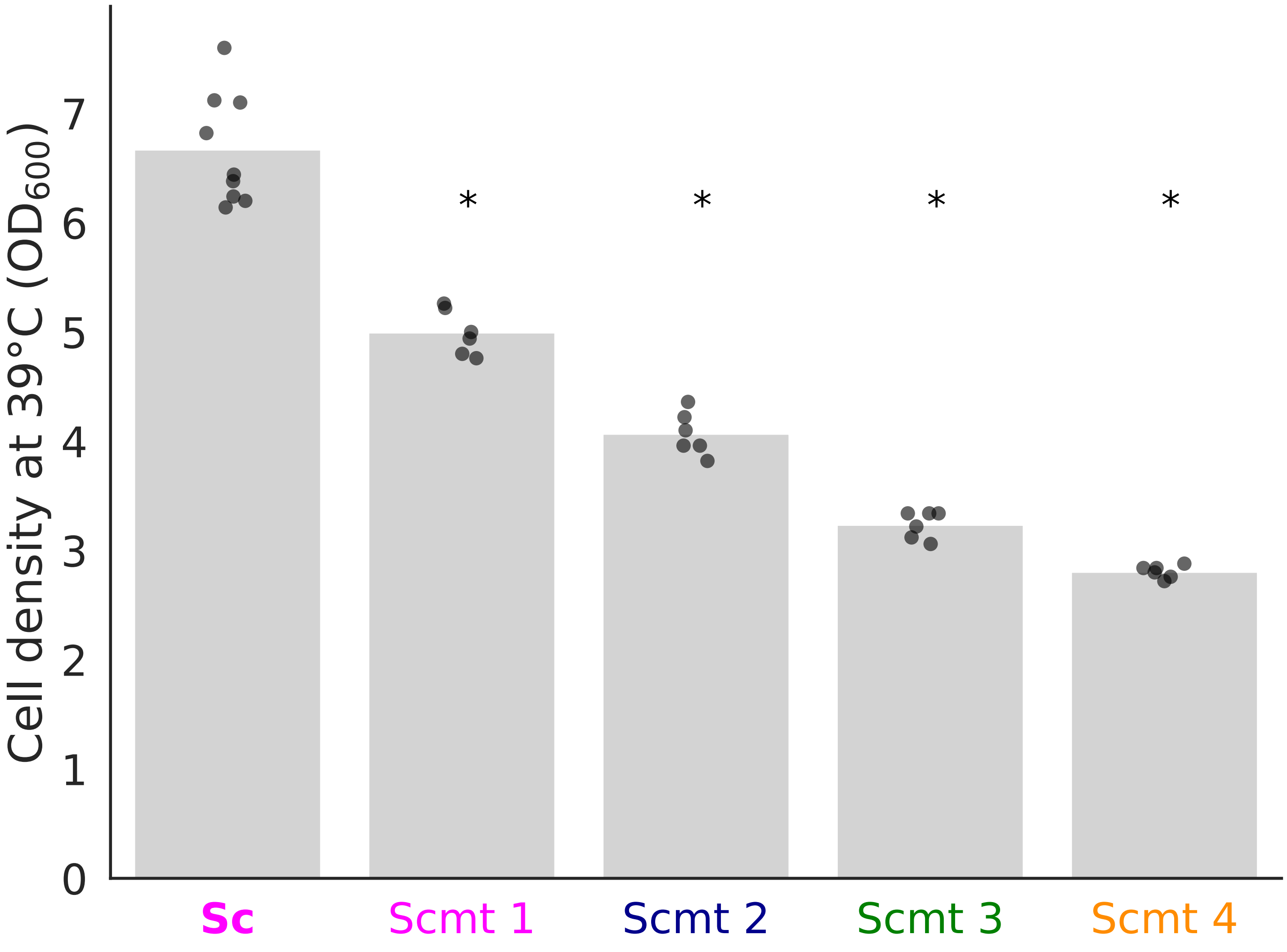

### Supplemental figure 7

Fig S7

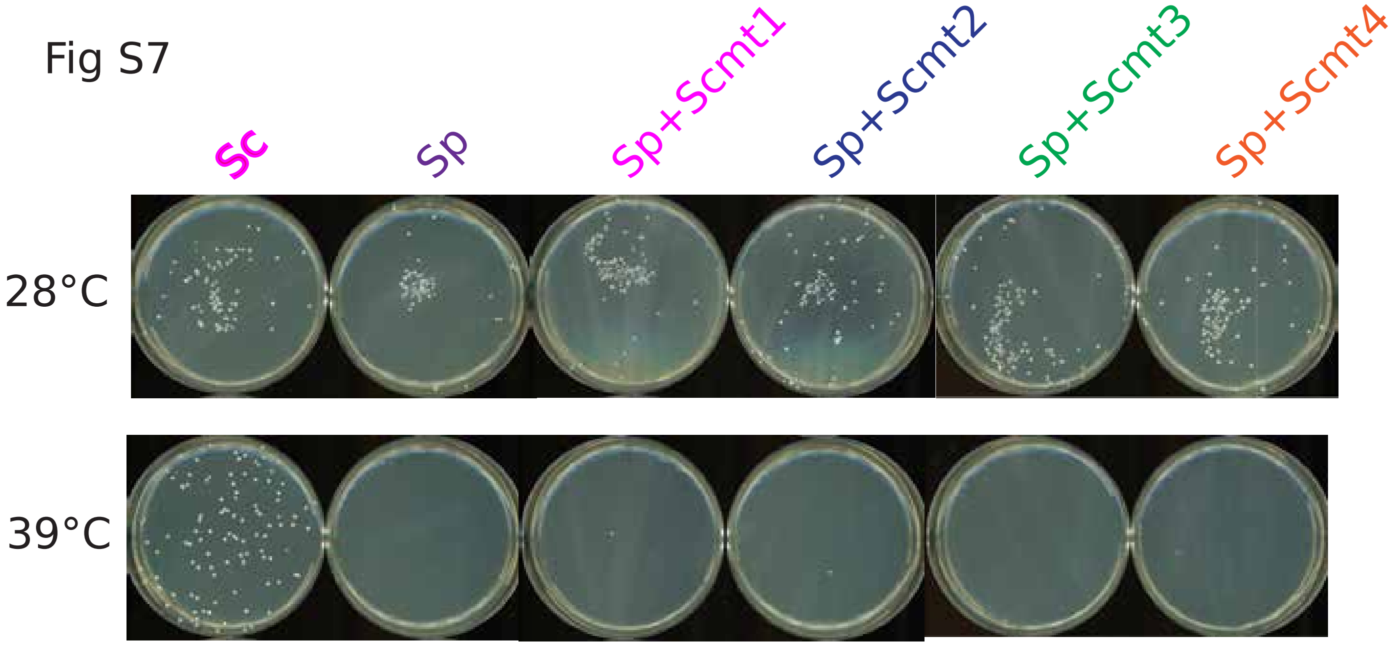

### Supplemental figure 8

Fig S8

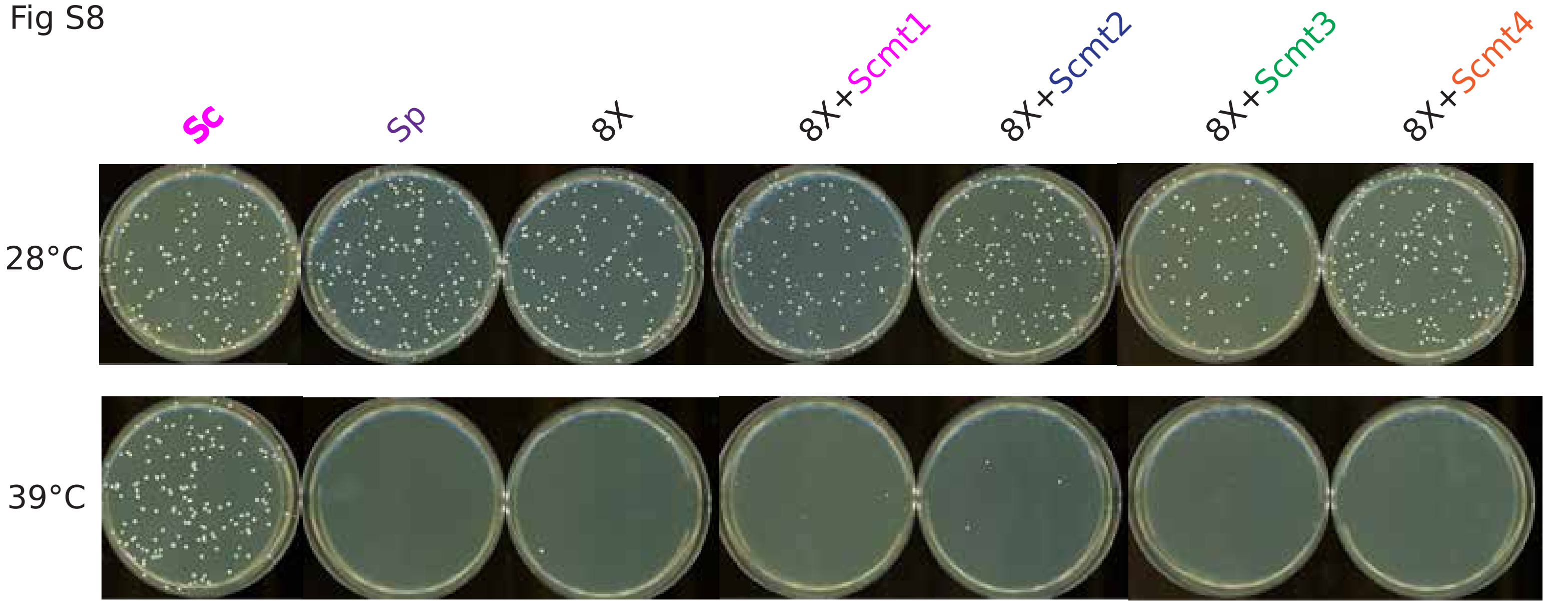

### Supplemental figure 9

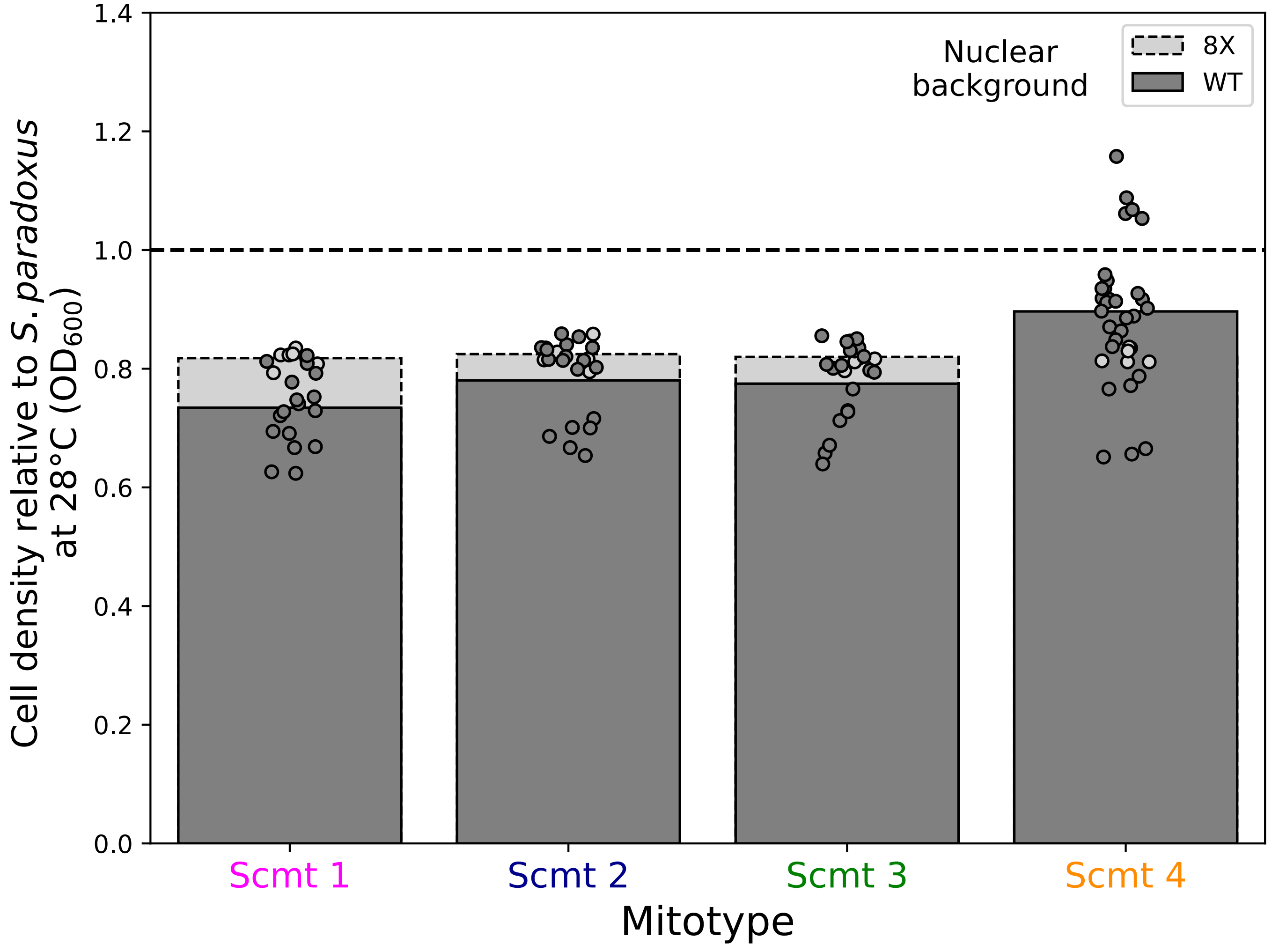

### Supplemental figure 10

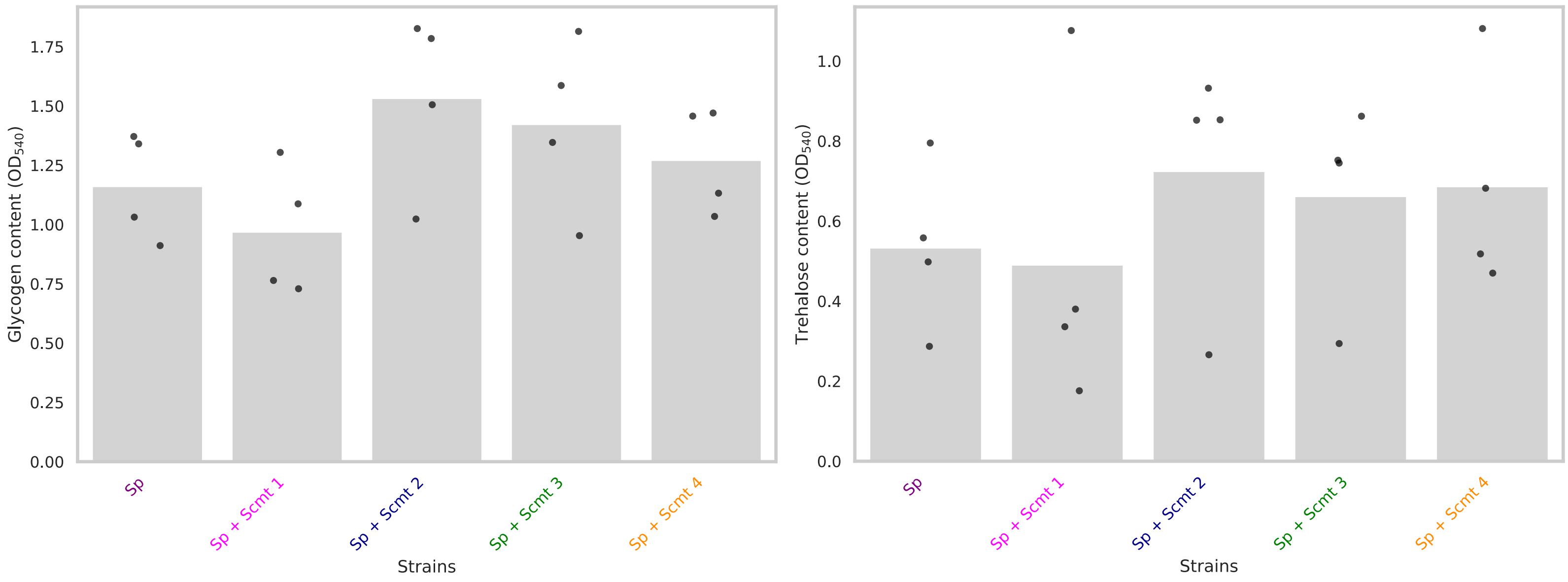

### Supplemental figure 11

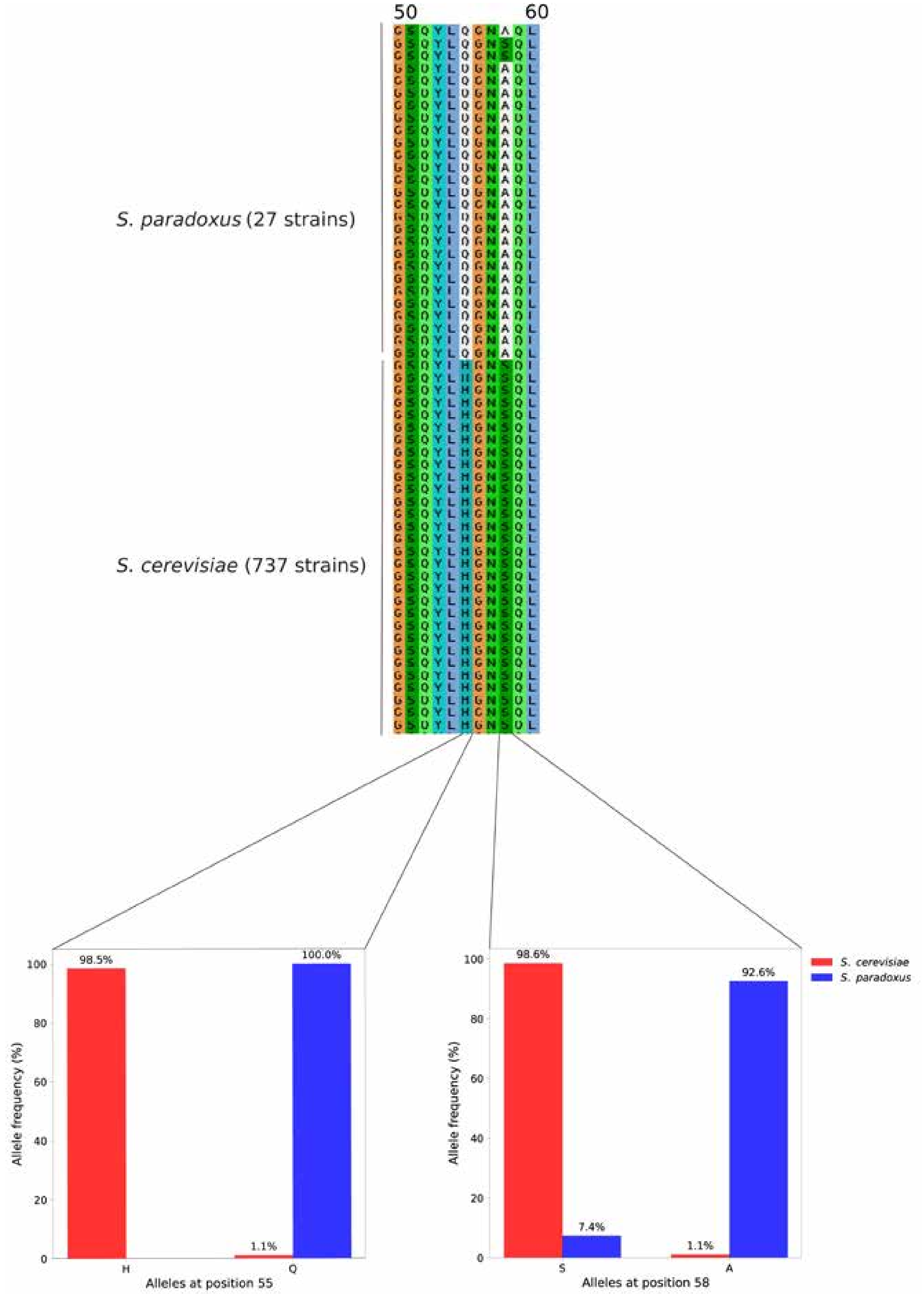
